# Introgression of type-IV glandular trichomes from *Solanum galapagense* to cultivated tomato reveals genetic complexity for the development of acylsugar-based insect resistance

**DOI:** 10.1101/2021.06.18.448858

**Authors:** Eloisa Vendemiatti, Rodrigo Therezan, Mateus H. Vicente, Maísa de Siqueira Pinto, Nick Bergau, Lina Yang, Walter Fernando Bernardi, Severino M. de Alencar, Agustin Zsögön, Alain Tissier, Vagner A. Benedito, Lázaro E. P. Peres

## Abstract

Glandular trichomes are involved in the production of food- and medicine-relevant chemicals in plants, besides being associated with pest resistance. In some wild *Solanum* species closely related to the cultivated tomato (*S. lycopersicum*), the presence of type-IV glandular trichomes leads to the production of high levels of insecticide acylsugars (AS). Conversely, low AS production observed in the cultivated tomato is attributed to its incapacity to develop type-IV trichomes in adult organs. Therefore, we hypothesized that cultivated tomatoes engineered to harbor type-IV trichomes on the leaves of mature plants can be pest resistant. We introgressed into the tomato cultivar Micro-Tom (MT) the capability of *S. galapagense* to maintain the development of type-IV trichomes throughout all plant stages, thus creating a line named “*Galapagos enhanced trichomes*” (MT-*Get)*. Mapping-by-sequencing of MT-*Get* revealed that five chromosomal regions of *S. galapagense* were present in MT-*Get*. Further mapping reveled that *S. galapagense* alleles on chromosomes 1, 2 and 3 are sufficient for the presence of type-IV trichomes, but in lower densities. GC-MS, LC-MS, and gene expression analyses demonstrated that the increased density of type-IV trichomes was not accompanied by high AS production and exudation in MT-*Get*. Moreover, MT-*Get* did not differ from MT in its susceptibility to whitefly (*Bemisia tabaci*). Our findings demonstrates that type-IV glandular trichome development along with AS production and exudation are partially uncoupled at the genetic level. The MT-*Get* genotype represents a valuable resource for further studies involving the biochemical manipulation of type-IV trichome content through either genetic introgression or transgenic approaches.

**Significance Statement:** This work identified loci in the tomato genome that control the heterochronic development of type-IV glandular trichomes and uncoupled the genetic control of this type of trichome ontogeny from acylsugar biosynthesis and accumulation, revealing a higher than anticipated genetic complexity of acylsugar-based insect resistance. The findings reported herein will contribute to further dissect the genetics of trichome development in tomato as well as to introgress broad and durable insect resistance in tomato and other Solanaceae.

## INTRODUCTION

Glandular trichomes have attracted considerable attention due to their economic potential as sources of a vast array of specialized metabolites (Schimiller *et al*., 2008; Tissier, 2012). Among such metabolites, many have industrial or medicinal value (Aharoni *et al*., 2005; Maes *et al*., 2011), whereas others are especially relevant in the protection against insect pests (Kang *et al*., 2010; Glas *et al*., 2012). The cultivated tomato and its wild relatives display great variation in trichome type, size, and number. Eight morphologically distinct types were defined for the *Lycopersicon* clade (the cultivated tomato and its 16 closest wild relatives), of which four are glandular: the types I, IV, VI, and VII (Luckwill, 1943; Glas *et al*., 2012). Type-IV trichomes (and, to a lesser extent, type-I trichomes, which are rare on tomato leaves) are sources of specialized metabolites called acylsugars (AS) (Kim *et al*., 2012; Ghosh *et al*., 2014; Ning *et al*., 2015; Schilmiller *et al*., 2015; Fan *et al*., 2016).

AS molecules consist of aliphatic acyl groups of variable chain lengths (C2 to C12) esterified to a glucose (G) or sucrose (S) moiety at four possible positions (Schilmiller *et al*., 2012; Ning *et al*., 2015; Fan *et al*., 2019). For instance, the AS molecule called S4:23 (2,4,5,12) is a sucrose-based AS esterified with C2, C4, C5, and C12 acyl groups, whose sum of aliphatic carbon atoms is 23. In the genus *Solanum*, AS confer resistance to fungal pathogens (Luu *et al*., 2017) and to multiple insect pests (Goffreda *et al*., 1990; Hawthorne *et al*., 1992; Liedl *et al*., 1995), such as whiteflies (*Bemisia spp*.), which is a major tomato (*S. lycopersicum*) pest worldwide (Maluf *et al*., 2010). AS deter insect and other arthropod attacks via distinct mechanisms, such as poisoning, sticking, and even “tagging” insects to increase predator recognition (Weinhold & Baldwin, 2011). Recently, it was demonstrated the horizontal transference of plant genes to whiteflies, which can be hijacked to neutralize plant-derived toxins (Xia *et al*., 2021). This kind of mechanism is unlike to evolve for AS detoxification, since these compounds act not only chemically but also mechanically to combat whiteflies. Therefore, AS comprise a robust and stable mechanism for whitefly resistance.

Type-IV trichomes have a single flat basal cell with a short two- or three-celled stalk (0.2-0.4 mm) and a round gland at the tip. They are particularly abundant in the wild species *S. galapagense, S. habrochaites*, and *S. pennellii* (Simmons & Gurr, 2005). These species produce much larger amounts of AS compared to the cultivated tomato (Fobes *et al*., 1985; Schilmiller *et al*., 2010). AS accumulation underlies the robust and multiple pest resistances of these wild tomato species (Mutschler *et al*., 1996; Momotaz *et al*., 2010; Rodriguez-Lopez *et al*., 2011; Leckie *et al*., 2012; Schilmiller *et al*., 2012). It was established that the cultivated tomato did not develop type-IV trichomes (Luckwill, 1943; Simmons & Gurr, 2005; McDowell *et al*., 2011; Glas *et al*., 2012). However, we have demonstrated that they do appear in the early stages of plant development (from the cotyledons to the 3^rd^ – 6^th^ leaf, depending on the cultivar), and that they can be used as markers of juvenility in tomato (Vendemiatti *et al*., 2017). Therefore, the lack of type-IV trichomes in the adult phase of the cultivated tomato explains the low presence of AS and susceptibility to herbivores.

Interestingly, the structural genes coding for the enzymes of the AS biosynthesis pathway are present in *S. lycopersicum* (Schilmiller *et al*., 2012; Schilmiller *et al*., 2015; Fan *et al*., 2016; Smeda *et al*., 2018). Indeed, four acylsugar acyltransferases (*ASAT1*–*ASAT4*) belonging to the BAHD acyltransferase superfamily were identified in the genomes of the cultivated tomato as well as its wild relatives (Schilmiller *et al*., 2012; Schilmiller *et al*., 2015; Fan *et al*., 2016; Fan *et al*., 2019). Biochemical characterization studies revealed that these *ASATs* sequentially catalyze the esterification of acyl chains at different positions of a sucrose moiety to generate a tetra-acylsucrose. Therefore, the catalytic activities of *ASATs* with different substrates can explain the diverse structures of *Solanum* spp. AS (Ghosh *et al*., 2014; Fan *et al*., 2016; Nadakuduti *et al*., 2017). Altogether, these observations led to the hypothesis that a cultivated tomato line modified to harbor type-IV trichomes on its adult leaves would accumulate high AS levels and be naturally resistant to pests.

Here, we successfully introgressed the capability of producing type-IV trichomes on adult leaves from *S. galapagense* LA1401 into the tomato genetic model system, cv. Micro-Tom (MT) (Carvalho *et al*., 2011). The introgressed MT line was named “*Galapagos enhanced trichomes*” (MT-*Get*) and showed a high density of type-IV trichomes on all leaves throughout plant development. The *S. galapagense*’s regions introgressed into MT-*Get* were determined by mapping-by-sequencing. This unique genetic material allowed us to determine the functionality of type-IV trichomes on cultivated tomato and its impact on insect resistance. For this end, we perfomed qRT-PCR analysis of *ASAT1-ASAT4* genes in leaves and the expression of *pSlAT2::GFP* in type-IV trichome glands. The AS profile of MT-*Get* was also determined by LC-MS and GC-MS. A preliminary assay of insect resistance was performed using *Bemisa* spp, one of the main insect pests of tomato and that is a target of AS. Our findings pave the way for molecular breeding of commercial varieties harboring a high density of type-IV trichomes and reveled the steps necessary to pursue insect resistance in the cultivated tomato.

## RESULTS

### 1. Introgression of the capacity to develop type-IV trichomes on adult leaves from *Solanum galapagense* LA1401 into tomato (*S. lycopersicum* cv. Micro-Tom)

*Solanum galapagense* LA1401 was chosen as a source of type-IV trichomes for genetic introgression. This accession is closely related to the cultivated tomato and we noticed that, unlike the cultivated tomato, it has a high density of type-IV trichomes on adult leaves, especially on the abaxial leaf surface (Figure **1a,b,d**). We have previously shown that the cultivated tomato plant produces type-IV trichomes only on cotyledons and juvenile leaves but not on adult leaves (Vendemiatti *et al*., 2017). We, therefore, set out to introduce the genetic determinants controlling the capacity to bear type-IV trichomes on adult leaves from *S. galapagense* into the cultivated tomato.

**Figure 1.**
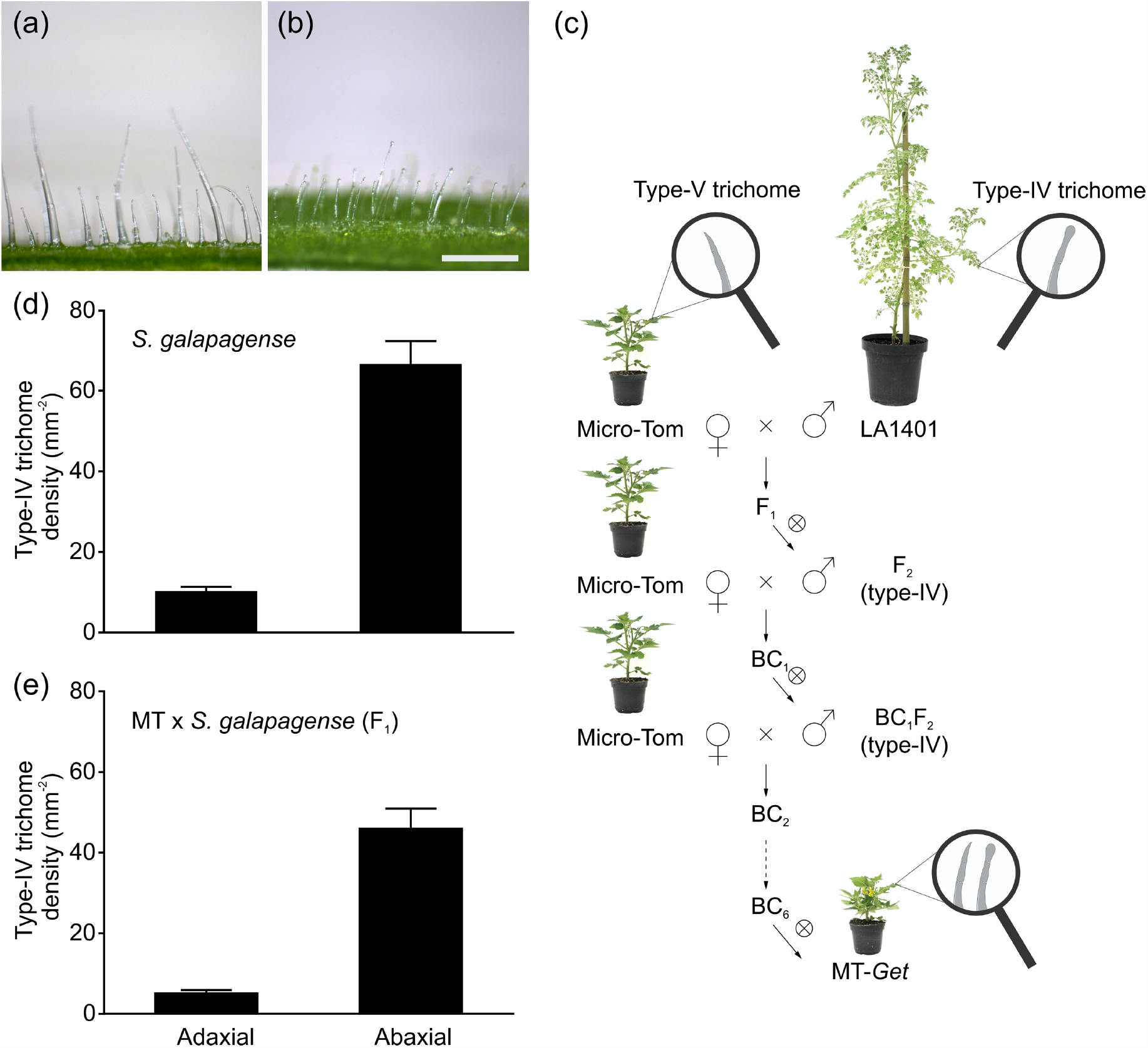
Introgression of *Solanum galapagense* (LA1401) type-IV trichome into the Micro-Tom (MT) model system. Representative light microscopy showing type-IV glandular trichomes on the adaxial (a) and abaxial (b) sides of the leaf. Scale bar=250 μm. (c) Introgression scheme used to create the Micro-Tom (MT) line bearing type-IV trichomes on adult leaves. The line was designated “*Galapagos enhanced trichomes*” (*Get*). X inside a circumference = self-pollination, BC= backcross. (d, e) Density (mm^- 2^) of type-IV trichomes on both leaf surfaces of the wild species (n=35) (d) and F_1_ plants (MT x *S. galapagense* LA1401) (n=30) (e). Data are mean ± SEM.

*Solanum lycopersicum* cv. Micro-Tom was fertilized with pollen from *S. galapagense*. After self-fertilization of F_1_ plants, we selected F_2_ plants with type-IV trichomes on leaves of developed plants. These plants were backcrossed (BC_1_) using MT as the recurrent parent. The process was repeated five more times until a stable BC_6_F_n_line that no longer segregated for the trait was obtained. The introgression scheme is shown in Figure **1c**. This new line was called “*Galapagos enhanced trichomes*” (MT-*Get*).

During the trichome characterization of F_1_ plants, we observed a lower density of type-IV trichomes on both leaf surfaces compared to the parental *S. galapagense*, (Figure **1d,e**), suggesting that this trait is dominant or semi-dominant with quantitative components.

We confirmed the identity of type-IV trichomes on adult leaves of MT-*Get* using scanning electron microscopy (Figure **2**). The type-IV trichome is a structure up to 0.4 mm tall with a glandular cell at the tip, and a unicellular and flat base (Luckwill, 1943; Channarayappa *et al*., 1992; Glas *et al*., 2012). This description fits with the structures shown in Figure **2c**. Thus, these results confirmed the presence of type-IV trichomes on adult leaves of MT-*Get*, which are similar in morphology and size to those present in *S. galapagense* (Figure **2d**).

We next verified whether the type-IV trichomes present in MT-*Get* were capable of expressing the acylsugar biosynthesis pathway. Transgenic MT and MT-*Get* plants harboring the *GFP* gene under the control of type-IV/I-specific *SlAT2* promoter (Schilmiller *et al*., 2012) were generated. Both MT and MT-*Get* cotyledons, as well as MT-*Get* adult leaves, displayed type-IV trichomes expressing GFP (Figure **3**). Accordingly, the absence of visible GFP signal on adult leaves correlated with the absence of type-IV trichomes (Figure **3f**). No GFP signal was detected in non-transgenic type-IV trichomes on the leaves of the MT-*Get* control (Figure **S2a,b**).

**Figure 2.**
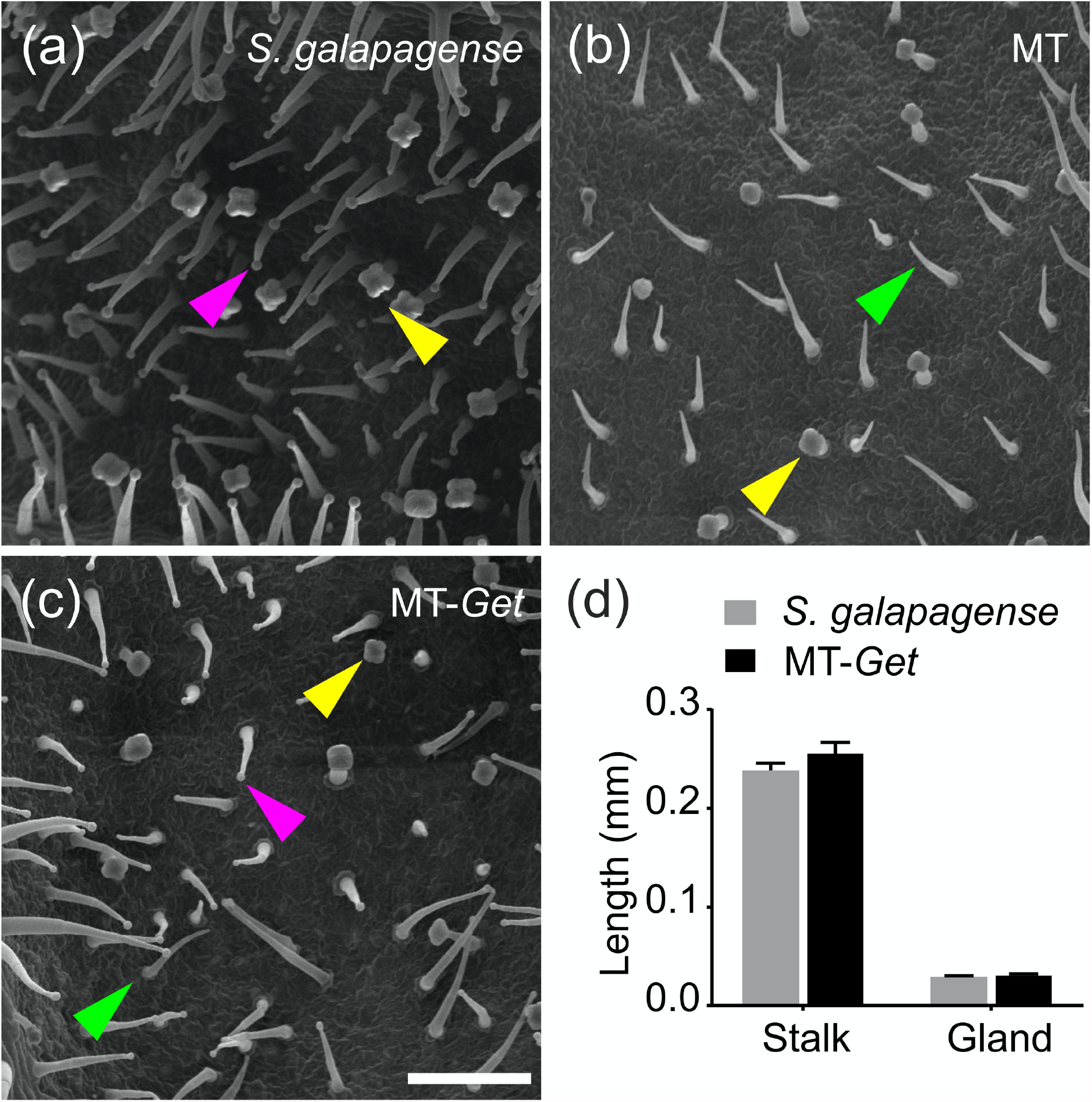
Scanning electron micrographs of abaxial surfaces of the 5^th^ leaf of representative 25-day-old plants of the tomato wild relative *Solanum galapagense* LA1401 (a), Micro-Tom (b), and the “*Galapagos enhanced trichomes*” line (MT-*Get*) (c). Scale bar=200 µm. The arrowheads represent the different trichomes: type IV (pink), type V (green), and type VI (yellow). (d) Type-IV trichome stalk height and gland size comparisons between *S. galapagense* and MT-*Get*. Data are mean (n=30) ± SEM. The data are not statistically different according to Student’s *t*-test (*P*< 0.05).

**Figure 3.**
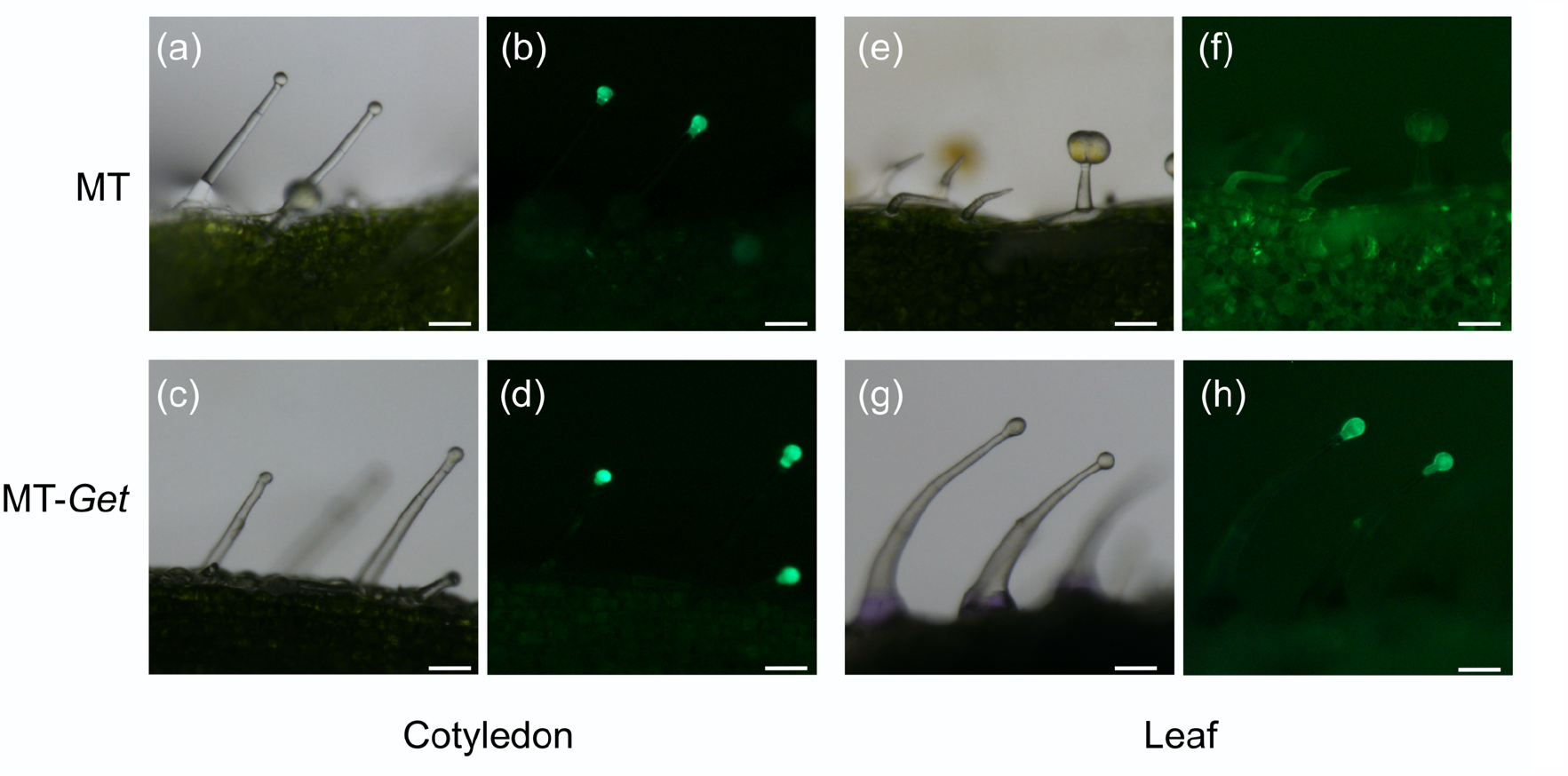
p*SlAT2::GFP* expression (green fluorescence) at the tip cells of MT and MT-*Get* type-IV trichomes on: the cotyledons (a juvenile organ) from MT (a, b) and MT-*Get* (c, d); and on the 5^th^ leaf (an adult organ) from MT (e, f) and MT-*Get* (g, h). Scale bar= 100µm.

### 2. Phenotypic characterization and genetic mapping of the “*Galapagos enhanced trichomes*” (*Get*) introgression line

The leaf developmental sequence of MT tomato from bottom to the top consists of a pair of embryonic leaves (cotyledons), a pair of juvenile leaves (1^st^ and 2^nd^ true leaves) and adult leaves (3^rd^ to 6^th^ upward) (Vendemiatti *et al*., 2017). To further characterize MT-*Get* plants, we first determined trichome classes and their respective densities on adult (5^th^) leaves. When comparing MT-*Get* and the control MT, an inverse relationship between type-IV and -V trichomes on both leaf surfaces was observed (Figure **4**). This pattern of type-IV predominance in MT-*Get* as opposed to type-V density in MT, was found in both juvenile as well as adult leaves (Figure **S3**). A lack of type-IV trichomes on MT adult leaves was observed (Figure **S3a,b**) and, on the other hand, MT-*Get* tended to lack type-V trichomes on juvenile leaves (Figure **S3c,d**). This inverse relationship between trichomes types IV and V had already been observed for several tomato cultivars in our previous report (Vendemiatti *et al*., 2017).

**Figure 4.**
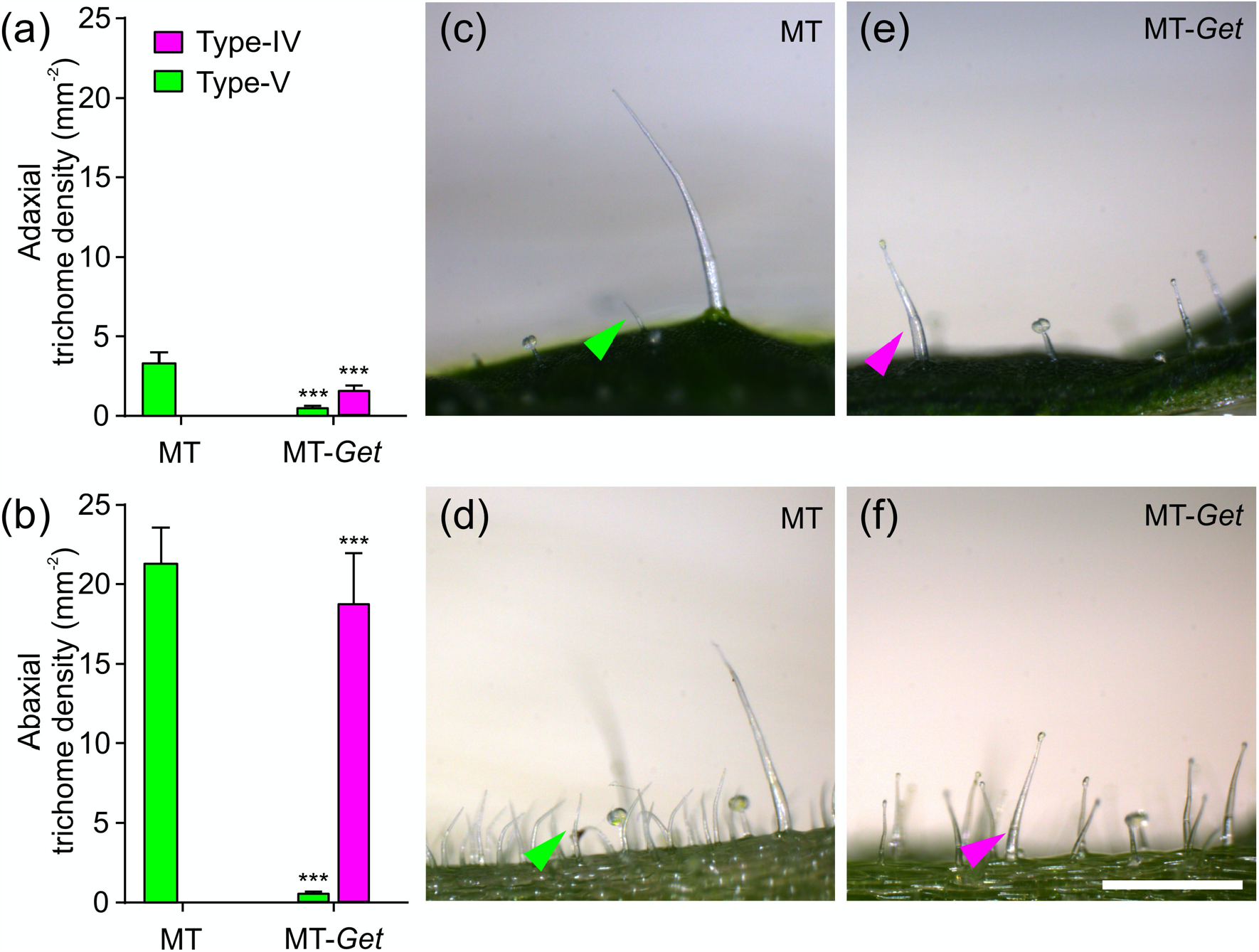
Density (mm^-2^) of types-IV and –V trichomes on the adaxial (a) and abaxial (b) surface of mature 5^th^ leaves of 45-day-old plants of Micro-Tom (MT) and MT-*Get*. (c, d) Representative micrographs of both surfaces of 5^th^ leave of 45-day-old MT plants. (e, f) Representative micrographs of both surfaces of 5^th^ leave of 45-day-old MT-*Get* (f) plants. Data are mean (n=40) ± SEM. Asterisks indicate a significant difference when compared with the reference sample according to the Student’s *t*-test at *P*< 0.001 (^***^). Scale bars= 250 μm.

When compared to *S. galapagense*, MT-*Get* displayed around 3.5-fold (abaxial) and 6.7-fold (adaxial) less type-IV trichomes (Figure **1d**; Figure **4**). *S. galapagense*, with only trichomes types I, IV and VI (Figure **1d**; **S4a**), showed less trichome diversity than MT-*Get*. Conversely, MT-*Get* bore trichomes types I, III, VI and VII (Figure **S4b, c**) in addition to the types IV and V (Figure **4**).

To determine the genetic configuration of MT-*Get, i*.*e*. the *S. galapagense* genome regions and alleles that were introgressed, we resorted to mapping-by-sequencing (Garcia *et al*., 2016). This approach allows to bulk-sequence the genomes of phenotypical categories of a segregating population to identify common loci responsible for a trait through the identification of distinct allelic frequencies between groups. MT-*Get* has a complex genetic composition: discrete *S. galapagense* genome segments were found on the long arms of MT chromosomes 1, 2 and 3, the short arm of chromosome 5, and a large pericentromeric region of chromosome 6 (Figure **5**). The genomic coordinates of the genetic variation from *S. galapagense* present at high frequencies (≥0.8) in the *Get*-like phenotypical group of the MT-*Get* segregating population are provided in Supporting Information Table **S2**. All the regions present on chromosomes 1, 2, 3, 5, and 6, or smaller combinations thereof, may be involved in type-IV trichome formation. Next, we segregated each fragment in sub-linages of *Get* using CAPS markers (Supplemental Table **S1**) that cover the extension of the fragments from *S. galapagense* in population derived from the Mapping-by-Sequencing experiment. We identified three sub-lines of MT-*Get* for the chromosomes 1, 2 and 3. Preliminary results revealed that these fragments are indeed involved with the type-IV trichome developmental pathway (Figure **S5**). When these three genomic fragments are isolated (Figure **S5c-h**), the type-IV trichome density is lower than in MT-*Get*, which carries all fragments (Figure **S5a,b**). These sequencing results suggest that the *Get* trait has a polygenic basis, involving epistatic interactions among multiple genes located in several genomic segments derived from *S. galapagense*, which probably control both trichome presence and density.

**Figure 5.**
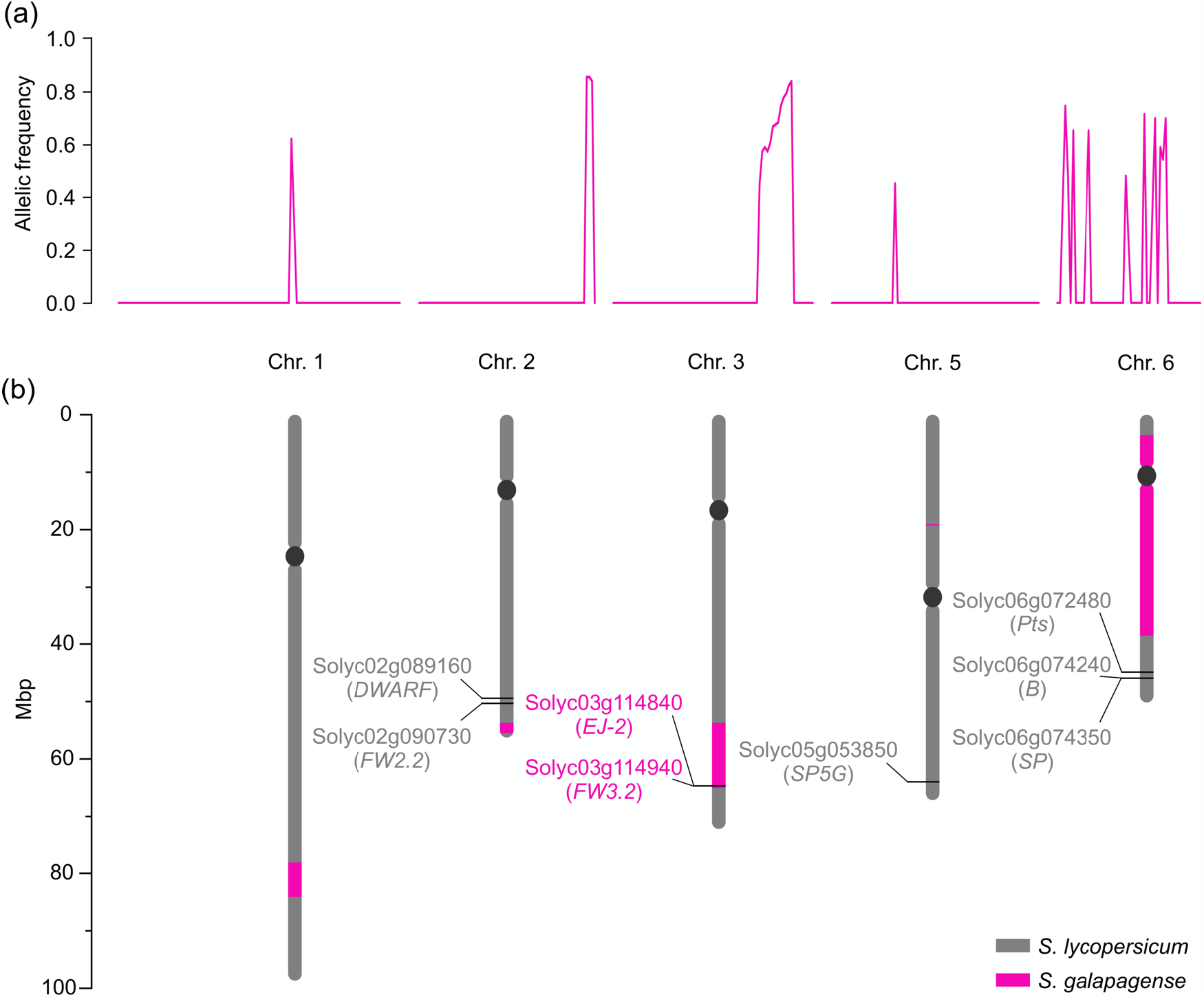
(a) Allelic frequency indicating the chromosomal positions where MT-*Get* has introgressions from *S. galapagense* LA1401. (b) Representation of the corresponding positions of the chromosomal fragments from *S. galapagense* (pink bars) introgressed into MT-*Get*. The positions were based on the *Solanum lycopersicum* cv. Heinz reference genome sequence. Relevant genes (as discussed in the text) and their SGN (Solyc) ID numbers are represented. Note that the MT-*Get* genome bears the MT-mutated alleles for *DWARF* (Bishop *et al*., 1996) and *SP* (Pnueli *et al*., 1998), which are determinant of the MT reduced plant size and determinate habit growth, respectively (Carvalho *et al*., 2011). The presence of the MT allele at the *SP5G* locus also contributes to the reduced plant size of MT-*Get*, since the *S. galapagense* allele promotes additional vegetative growth due to a lack of flower induction under long days (Soyk *et al*., 2017b). MT-*Get* also lacks *S. galapagense* alleles for genes conferring additional phenotypes distinct to this wild species, such as highly dissected leaves (*Pts*) (Kimura *et al*., 2008) and β-carotene accumulating fruits (*B*) (Ronen *et al*., 2000). The impact of the wild species alleles *EJ-2* and *FW3*.*2* on the MT-*Get* phenotype is shown in Figure S6 and discussed in the text.

We also verified whether the presence of known alleles from *S. galapagense* in the introgressed segments could contribute to additional, developmental differences independent of trichome traits by comparing the distinct genomic regions between MT-*Get* and MT. Notably, within the chromosome 3 segment, MT-*Get* harbors the *S. galapagense* alleles for the genes *EJ-2* (Soyk *et al*., 2017a) and *FW3*.*2* (Chakrabarti *et al*., 2013) (Figure **5b**). The pleiotropic effects of non-domesticated alleles of the *FW3*.*2* gene, which codes for a *P450 monooxygenase*, are known to lead to a reduction in fruit weight and increased shoot branching (Chakrabarti *et al*., 2013). Both phenotypes are present in MT-*Get* (Figure **S6a-d**), which harbours the *S. galapagense fw3*.*2* allele. Another gene controlling fruit weight is *FW2*.*2* (Frary *et al*., 2000), although it cannot be responsible for the smaller fruit of MT-*Get* compared to MT because both genotypes harbour the same MT allele (Figure **5b**). Despite the large chromosome 6 segment from *S. galapagense* introgressed into MT-*Get* (Figure **5b**), this line has the same red fruit characteristic of MT (Figure **S6e,f**). This is due to the absence of the *S. galapagense B* allele, which is responsible for orange fruits (Figure **S6g**) and also maps on the long arm of chromosome 6 (Ronen *et al*., 2000) but, accordingly, outside the introgressed region. Lastly, the reduced size of MT-*Get* sepal (Figure **S6e,f**) is probably an effect of the *EJ-2* allele from *S. galapagense* (Figure **S6h**), since this *MADS-box* gene controls the size of organ in this floral whorl (Soyk *et al*., 2017a).

### 4. Whitefly resistance test and trichome exudation in MT-*Get* plants

Based on the fact that type-IV trichomes drive whitefly (*Bemisia tabaci*) resistance in the same accession of *S. galapagense* used here (Firdaus *et al*., 2012; Firdaus *et al*., 2013; Vosman *et al*., 2019), we verified whether MT-*Get* displayed an increased resistance to this insect. However, MT-*Get* did not differ from MT (Figure **S7a-c**) in a preliminary assay based on the number of whitefly nymphs on leaves after exposure to a controlled greenhouse infested with whiteflies. We also observed that MT-*Get* did not display exudates at the tip of the type-IV trichome gland. The production of such exudates which is a feature typical of *S. galapagense* (Figure **S7d,e**) that accounts for its sticky leaves and is regarded to be the effect of AS accumulation (Schilmiller *et al*., 2008). We also observed that, differently from *S. galapagense*, MT-*Get* leaves were not sticky to the touch. *S. galapagense* exudates can be stained with Rhodamine-B, which is a dye for AS. In MT-*Get*, Rhodamine-B staining was restricted to the area inside the gland (see inserts in Figure **S7d,e**). These results prompted us to profile the AS accumulated on leaves and the expression of type-IV trichome-specific AS biosynthesis genes (Schilmiller *et al*., 2012; Fan *et al*., 2015; Schilmiller *et al*., 2015) in MT-*Get*.

### 5. Acylsugar accumulation and related gene expression in MT-*Get*

Since type-IV trichomes are the main sources of AS (Goffreda *et al*., 1989; Liedl *et al*., 1995; Maluf *et al*., 2010), we hypothesized that this insecticide would accumulate on adult leaves of the cultivated tomato upon the introduction of the capacity to maintain the development of type-IV trichomes. We, therefore, conducted a metabolic profile analysis using both liquid chromatography and mass spectrometry to assess AS accumulation on adult leaves of the MT-*Get*, compared to the parental *S. galapagense*.

*S. galapagense* showed peaks corresponding to a variety of sucrose (S)-based AS with different acyl moieties, ranging from 2 to 12 carbons (C2 to C12) (Figure **6**, Table **1**). Consistently, the AS peaks were very attenuated or absent in the cultivated tomato (MT), which is already known to accumulate very low amounts of AS (Figure **6**) (Blauth *et al*.,1998). MT-*Get*, although accumulating more AS than MT, showed dramatic quantitative and qualitative differences when compared to *S. galapagense*. Notably, MT-*Get* showed reduced levels of AS harboring C10 and C12 moieties, such as S4:23 (2,4,5,12), S4:22 (2,5,5,10), and S4:24 (2,5,5,12) (Figure **6**, Table **1**). The amounts of S4:23, S4:22, and S4:24 were 120, 42, and 18-fold lower in MT-*Get* than *S. galapagense*, respectively (Table **1**). These differences are greater than those for type-IV trichome densities between *S. galapagense* and MT-*Get*, which were 3.5-fold (abaxial surface) and 6.7-fold (adaxial surface) (Figure **1**, **4**).

**Figure 6.**
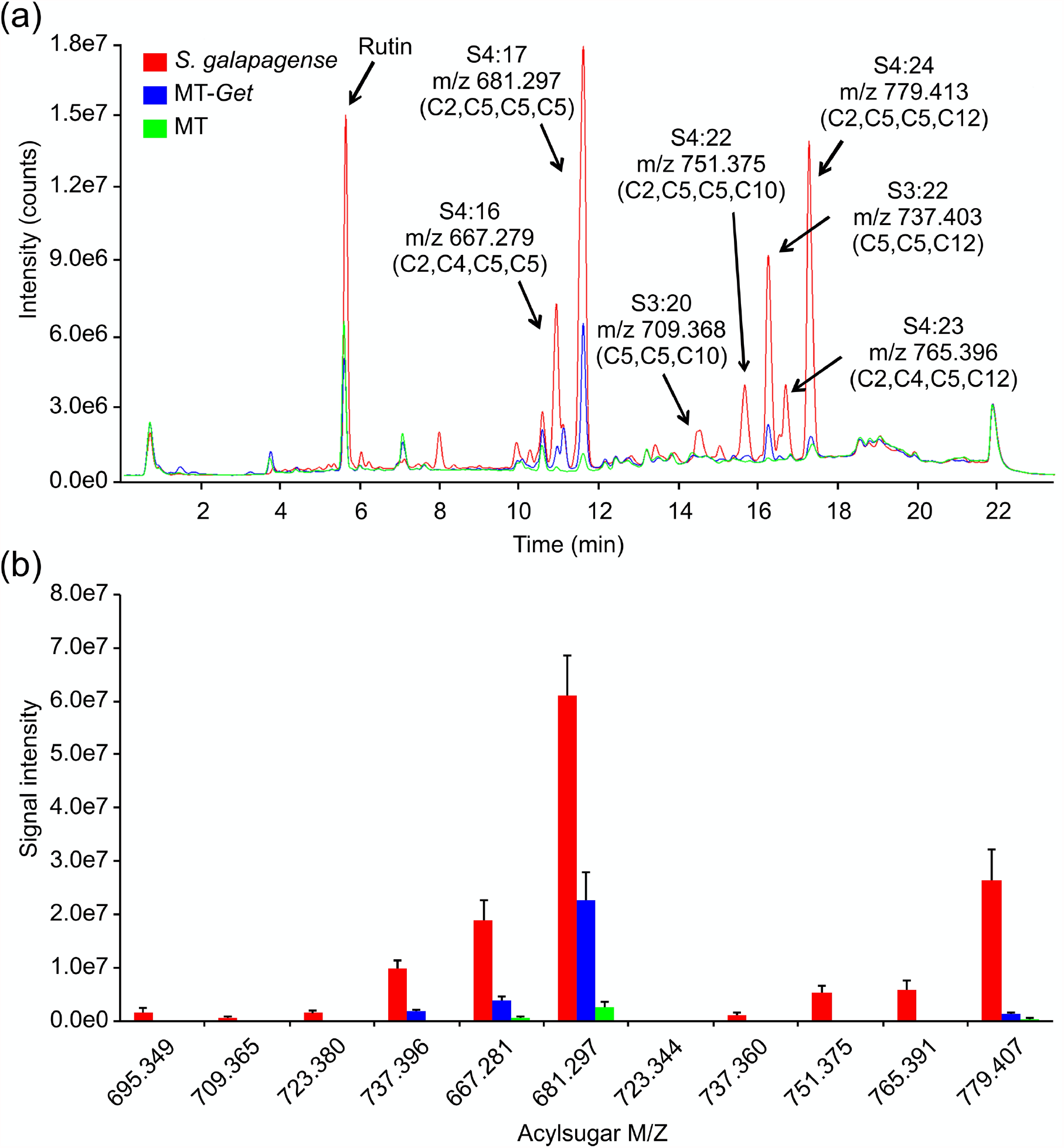
Acylsugar (AS) content in *Solanum galapagense* LA1401, Micro-Tom (MT), and MT-*Get*. (a) Representative LC-MS chromatogram. Peak area quantifications are shown in Table 1. (b) Signal intensity for each one of the AS analyzed in the three genotypes.

**Table 1.** Peak areas from ion chromatograms (LC-MS) of acyl sugars from *S. galapagense*, MT-*Get* and MT.

| AS | *m/z* | Peak area |  |  | *S.galap./*<br>MT-*Get* |
| --- | --- | --- | --- | --- | --- |
|  |  | *S. galapagense* | MT-*Get* | MT |  |
| S3:20 (5,5,10) | 709.368 | 675.914 **a** | 72.873 **b** | 7.192 **c** | 9.28 |
| S3:22 (5,5,12) | 737.403 | 9,877.349 **a** | 1,767.007 **b** | 145.07 **c** | 5.59 |
| S4:16 (2,4,5,5) | 667.279 | 18,907.925 **a** | 3,835.183 **b** | 514.101 **c** | 4.93 |
| S4:17 (2,5,5,5) | 681.297 | 61,140.597 **a** | 22,592.369 **b** | 2,509.92 **c** | 2.71 |
| S4:22 (2,5,5,10) | 751.375 | 5,363.138 **a** | 125.538 **b** | 26.872 **c** | 42.72 |
| S4:23 (2,4,5,12) | 765.396 | 5,864.023 **a** | 48.632 **b** | 26.809 **c** | 120.58 |
| S4:24 (2,5,5,12) | 779.413 | 26,335.227 **a** | 1,446.972 **b** | 301.079 **c** | 18.20 |

Acyl sugars were identified according to their *m/z* and retention time (n=4). For the sake of simplicity values were divided by 1000. Values followed by different letters in each row are statistically different according to Student's *t*-test ( $P < 0.05$ ).

In the GC-MS analysis, detectable peaks of n-decanoate (C10) and n-dodecanoate (C12) were observed only for *S. galapagense* (Figure **S8**, Table **S3**). These carboxylates are derived from C10 and C12 harboring acylsugars, which agrees with the higher amounts of the acylsugars S4:23 (2,4,5,12), S4:22 (2,5,5,10) and S4:24 (2,5,5,12) found in the LC-MS analysis in *S. galapagense* (Figure **6**). Only small quantities of methyl dodecanoate were detected in MT-*Get* (Table **2**), which correlates with the presence of S3:22 (5,5,12) and S4:24 (2,5,5,12) in this genotype (Figure **6**).

We next evaluated the relative expression of the know genes involved in AS biosynthesis. There are four acyltransferases (ASAT) that act sequentially to esterify acyl chains in specific positions of the sugar moiety (Schilmiller *et al*., 2012; Fan *et al*., 2016). The expression levels of the four *ASAT* genes were higher in *S. galapagense* leaves compared to MT-*Get* (Figure **7**), which correlates with the differences in AS content determined (Figure **6**, Table **1**).

**Figure 7.**
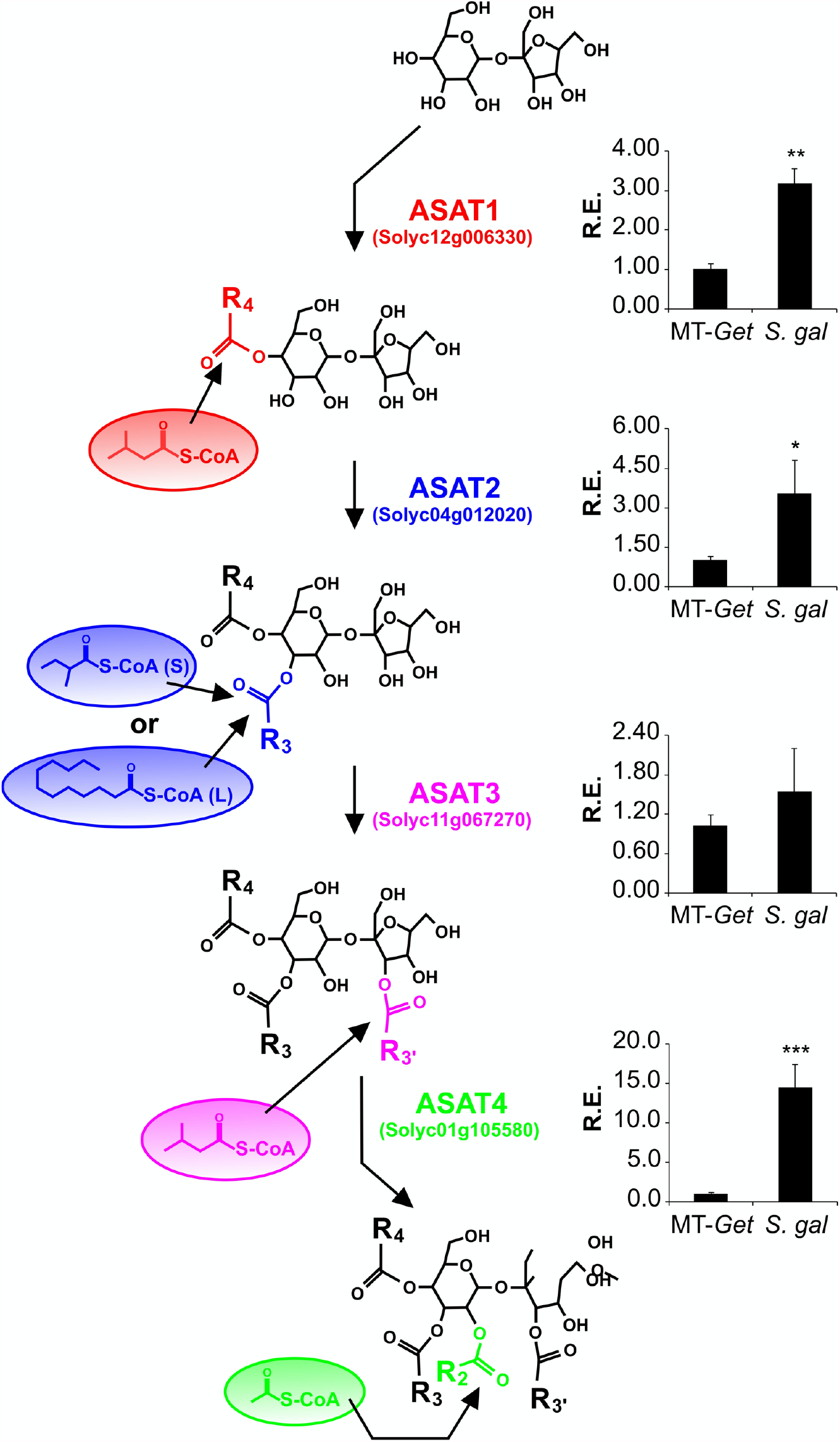
Relative transcript accumulation of acylsugar acyltransferase (*ASAT*) genes in leaf tissues of MT-*Get* and *S. galapagense*. qRT-PCR values are means ± SE. Asterisks indicate statistically significant differences when compared with the reference sample according to the Student’s *t*-test at *P*< 0.05 (^*^); *P*< 0.01 (^**^); *P*< 0.001 (^***^).

The relative expression of the genes coding for acylhydrolase (ASH) enzymes was also quantified (Figure **S9a-c**). They are responsible for the removal of acyl chains from specific AS positions, thus creating the substrate for the action of ASAT (Schilmiller *et al*., 2016; Fan *et al*., 2019). The relative expression of *ASH*1 in MT-*Get* was significantly higher compared to *S. galapagense* (Figure **S9a**), which might also reflect the differences in AS content observed between these two genotypes.

Since MT-*Get* and *S. galapagense* seem to differ in the capacity to exudate AS (Figure **S7**), we assessed the gene expression of a putative efflux transporter, which may be responsible for AS exudation in type-IV trichome tips. Notably, the ABC transporter (Solyc03g005860) previously associated with AS exudation (Mandal *et al*., 2019) presented a higher expression level in *S. galapagense* compared to MT*-Get* (Figure **S9d**).

## DISCUSSION

Type-IV trichomes are involved in important mechanisms for herbivore resistance in the *Solanum* genus and beyond. In the wild species *S. pennellii, S. galapagense*, and *S. habrochaites*, these structures are found at high densities, making them AS accumulators (Simmons & Gurr, 2005; Mutschler *et al*., 1996; Momotaz *et al*., 2010; Leckie *et al*., 2012; Schilmiller *et al*., 2012). These specialized metabolites protect plants via both their toxicity and their stickiness, thereby trapping and immobilizing the insects, or labeling them for predator recognition (Mirnezhad *et al*., 2010; Weinhold & Baldwin, 2011; Vosman *et al*., 2019; Schuurink & Tissier, 2019). Since type-IV trichomes are not found on the adult structures of the cultivated tomato (*S. lycopersicum*) (Vendemiatti *et al*., 2017), obtaining and studying a line with this phenotype is a critical step towards a better understanding of broad insect resistance based on acylsugars as well as elucidating the molecular mechanisms of glandular trichome development. We created this line by introgressing into *S. lycopersicum* cv. Micro-Tom the trait from *S. galapagense* LA1401, which is a wild species related to the cultivated tomato and highly resistant to whiteflies (Firdaus *et al*., 2013). We named the introgression line “*Galapagos enhanced trichomes*” (MT-*Get*).

MT-*Get* displays several traits that overall look intermediate between both parents: it has less glandular trichomes than *S. galapagense*, but a higher diversity of trichome types, whereas the comparison with MT has the opposite trend: MT-*Get* has more glandular trichomes and less diversity (Figure **8a**). The reduction of the trichome diversity in *S. galapagense* and MT-*Get* is mainly represented by the reduced densities (or absence) of trichomes types III, V, and VI. In the case of type-V trichomes, its inverse correlation with type-IV structures had already been pointed out in a previous study comparing juvenile and adult leaves of cultivated tomato cultivars (Vendemiatti *et al*., 2017). This result suggests that both trichome types may have an overlapping ontogeny since they only differ anatomically by the presence/absence of a terminal gland (Luckwill, 1943; Glas *et al*., 2012). The density reduction of trichomes types III and VI might be also a consequence of the increased number of type-IV trichomes via a general mechanism of trichome initiation and differentiation that controls the identity of neighboring epidermal cells. The mechanism that prevents trichome formation in clusters is well known in Arabidopsis (Pesch & Hülskamp, 2011) and was recently also suggested for tomato (Schuurink & Tissier, 2019). Our findings reveal the complexity of studying trichome distribution as a trait in tomato since perturbations in one trichome type are likely to produce a pleiotropic effect in the abundance of other types.

**Figure 8.**
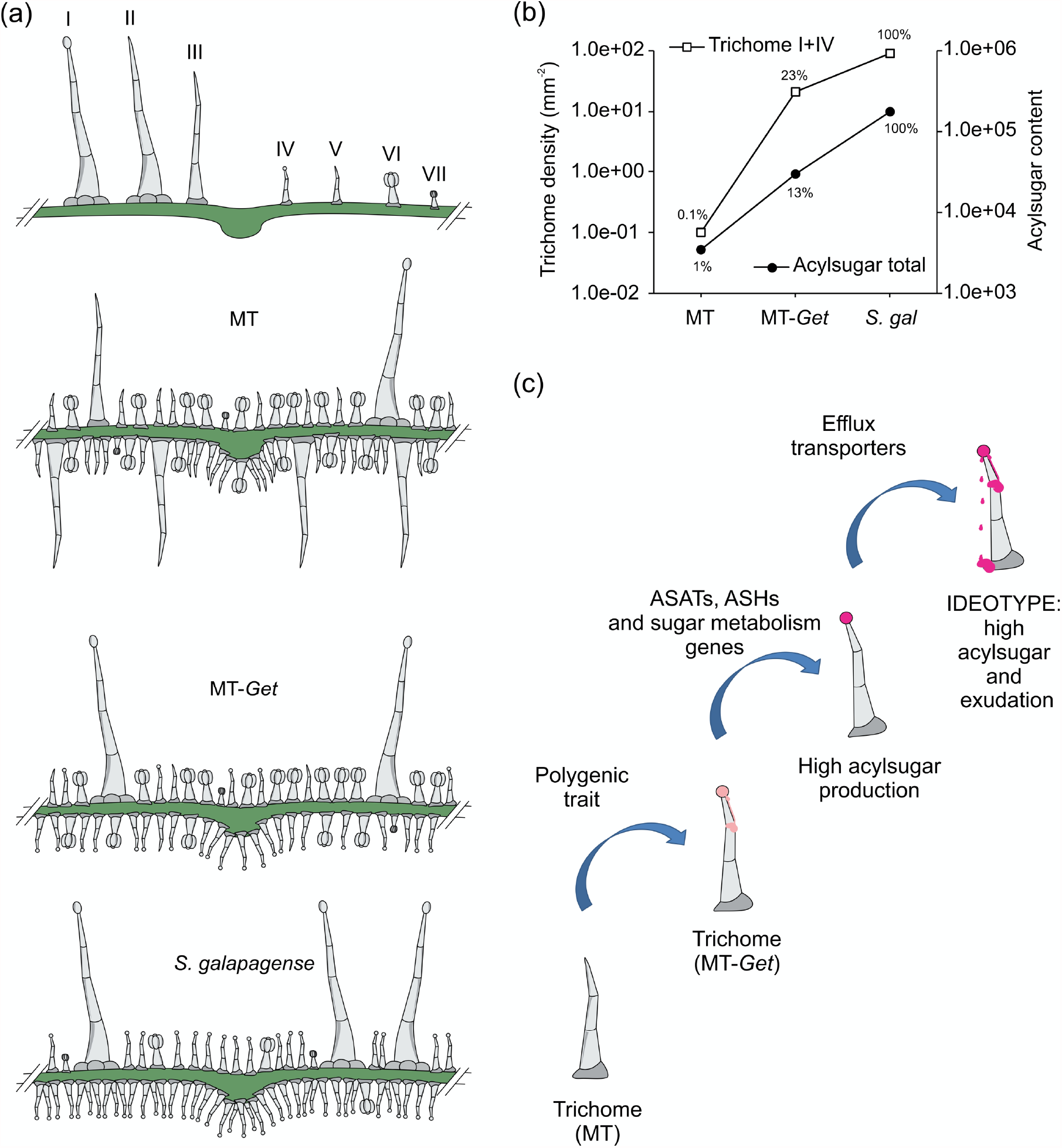
(a) Schematic representation of trichome type distribution in each genotype used in this work. (b) Logarithmic scale representation of the results of total trichomes related to the total acylsugar content (types I and IV). (c) Sequential steps to obtain broad and durable insect-resistant tomatoes through the high production of acylsugars on tomato leaves.

Genetically, our analysis revealed that the development of type-IV trichomes is associated with multiple genes dispersed over several chromosomes. It is not clear yet whether all genomic fragments (loci) identified in this study are essential for type-IV trichome development. However, this result led us to hypothesize that the trait has a polygenic inheritance and that some of the genes within these regions can explain the additional traits present in the introgressed lineage. One region found on chromosome 2 coincides with a previous QTL associating adult whitefly survival and the presence of type-IV trichomes in segregating populations (F_2_and F_3_) from a cross between the cultivated tomato and *S. galapagense* (Firdaus *et al*., 2013; Vosman *et al*., 2019). This is strong evidence that at least one gene that is necessary for type-IV trichome formation is located in this region of the genome. However, the preliminary analysis of sublines derived from MT-*Get* harboring *S. galapagense* alleles only in this region suggests that it is not sufficient for high type-IV densities. This same conclusion can be made for the *S. galapagense* alleles present on chromosome 1 and 3, whose corresponding sublines presented type-IV trichomes, but in lower densities. Therefore, it is likely that the high density of type-IV trichomes in MT-*Get* could be the result of epistatic interactions of different *S. galapagense* alleles. On chromosome 3, MT-*Get* also harbors *S. galapagense* alleles for two known developmental genes: *EJ-2* and *FW3-2*. However, it is unlikely that they can be directly associated with type-IV trichome development since these structures are absent in adult leaves of other species in the tomato clade carrying wild alleles of *EJ-2* and *FW3-2*. This is the case of the *S. cheesmaniae* accessions LA0521 and LA1139, and *S. pimpinellifolium* LA4645 (Simmons & Gurr, 2005; Bitew, 2018).

The initial hypothesis that the presence of type-IV trichome was sufficient for high AS production and herbivore resistance was not borne out by our data. Our results show that MT-*Get* plants, although having a reasonable density of type-IV trichomes, do not produce AS on the same scale as *S. galapagense* (Figure **8b**). In agreement with this, the MT-*Get* line did not show improved resistance to *Bemisia tabaci*, the main insect controlled by the presence of type-IV trichomes in the wild species (Maluf *et al*., 2010). One interesting feature observed here was the differences between MT-*Get* and *S. galapagense* in the accumulation of AS with medium-chain carbon groups (10 and 12 carbons). These results suggest that among the genes controlling the AS metabolic pathway in *S. galapagense*, some could be related to the esterification of C10 and C12 acyl groups, which may have repercussions for the level of insect resistance.

In our evaluation of the known structural genes of the AS pathway, we noticed that *ASAT1* (Solyc12g006330), *ASAT2* (Solyc04g012020), *ASAT3* (Solyc11g067270), and *ASAT4* (Solyc01g105580) are not located within any of the introgressed regions (See Figure **5**). This means that *S. galapagense* and MT-*Get* have distinct alleles for these genes, including potentially *cis*-regulatory elements. Therefore, the effect of both *cis* and *trans* elements on regulating the expression of *ASAT* genes may explain the higher expression of *S. galapagense* compared to MT-*Get*. However, we cannot exclude that the differences in transcript accumulation may also reflect in part the enriched content of type-IV trichome-derived RNAs in *S. galapagense* due to their higher density compared to that of MT-*Get* (Figure **1d**, **4**). On the other hand, the difference in *ASAT4* expression between MT-*Get* and *S. galapagense* is far beyond the magnitude of the trichome density difference between these genotypes. It is interesting to note that the higher expression of *ASAT2* in *S. galapagense* is consistent with the observation of increased levels of C10-12 acyl groups in the wild parental genotype (Figure **6**). We propose that the enzyme encoded by *ASAT2* from *S. galapagense* may be able to esterify more efficiently medium-acyl chains (up to 12 carbons) in the R3 position of the sucrose backbone than the MT allele (Fan *et al*., 2016; Figure **7**).

Since AS are non-volatile compounds, they are produced in the glands and by a mechanism that is not yet clear, drip out of the gland (Schuurink & Tissier, 2019). This phenomenon was observed for *S. galapagense* (Figure **S7d**) and may sustain the positive feedback responsible for AS production. A comparative transcriptomic analysis of *Solanum pennellii* accessions with distinct AS contents found that the expression levels of most AS metabolic genes were positively correlated with AS accumulation (Mandal *et al*., 2019). Among the differentially expressed genes (DEGs), three genes putatively encoding ATP-binding cassette (ABC) transporters were upregulated in the accessions with high AS content. Furthermore, Dimissie *et al*. (2019) described an ABC transporter strongly expressed in the glandular trichomes of *Lavandula angustifolia* (Lamiaceae). Based on this information, we verified the relative expression of an ABC transporter (Solyc03g005860) in our material and found the same pattern of *ASAT* expression, i.e., its gene expression was closely related to the type-IV trichome density on the leaves (Figure **S9d**). While this result suggests that this ABC transporter may be involved in AS exudation, the actual factors accounting for the differences in AS exudation capacity between *S. galapagense* and MT-*Get* remains to be discovered. It is worth noting that AS transport could be critical in determining how much AS is produced and secreted. One possibility is that AS would remain inside the trichome head in the absence of efficient transport, leading to feedback inhibition. On the other hand, efficient transport might drive biosynthesis by creating a metabolic flux, thereby potentially preventing feedback inhibition, and ultimately leading to high AS accumulation.

<u>W</u>e initially expected that type-IV trichomes of *S. galapagense* would have the capacity to accumulate the amounts of sugar moieties necessary to be acylated in the gland tip. Studies using radiolabeled carbon in tobacco showed that isolated trichome glands might be metabolically independent to produce the main exudates, but only when adequately supplied with carbon sources (Kandra & Wagner, 1988). Earlier transcription analyses performed with expressed sequence tags (EST) indicated that trichomes could work with simple biochemical input while having few highly active biochemical pathways of the primary and specialized metabolisms locally and highly active (Schilmiller *et al*., 2008). Although type-IV trichomes contain chloroplasts (Figure **S2c**), these probably are not in sufficient number to sustain the primary as well as the specialized metabolisms occurring in the cells of this structure (Schilmiller *et al*., 2008; Balcke *et al*., 2017). Therefore, the differences in AS accumulation are unlikely to be fully explained by genes related to modifications of the acyl moiety, such as *ASATs* and *ASHs*. Still, instead, there could be additional unknown genes involved in sugar metabolism or transport that can enable the trichome gland to become a stronger sink.

## Concluding remarks

The results presented here and their implications can be summarized in a model in which the transfer of insect resistance from a wild species into the cultivated tomato requires the stacking of three types of genetic determinants: i) Favourable alleles necessary to build the specific glandular trichomes at a correct developmental stage, such as in MT-*Get*; ii) Favourable alleles necessary for specific metabolic pathways (*e*.*g*. different compositions of acyl groups and capacity to accumulate the sugar moiety), and iii) Favourable alleles necessary to transform glandular trichomes into exudation structures, such as transmembrane transporters (Figure **8c**). The MT-*Get* introgression line presented here is the starting point of a challenging, long-sought breeding goal – the introduction of a trait in tomato for effective, broad, long-lasting insect resistance, and decrease the pesticide use. Altogether, this study demonstrates that glandular trichome development along with the metabolite production pathway and exudation are partially uncoupled at the genetic level. The MT-*Get* genotype represents a valuable resource for further studies involving the biochemical manipulation of type-IV trichome content through either genetic introgression or transgenic approaches. *In toto*, MT-*Get* is the first step to creating a tomato plant that naturally produces a substance that actively kills pests. In other words, we have created plants that carry the weapon, but we require a deeper understanding of genetics to load it with the appropriate metabolic ammunition.

## EXPERIMENTAL PROCEDURES

### Plant material, growth conditions and breeding scheme

Seeds of *Solanum galapagense* LA1401 were obtained by the Tomato Genetics Resource Center (TGRC - University of California). Micro-Tom (MT) seeds were donated by Dr. Avram Levy (Weizmann Institute of Science, Israel) and maintained through self-pollination as a true-to-type cultivar since 1998. The “*Galapagos enhanced trichome*” (MT-*Get*) line was generated by the cross MT x *S. galapagense* LA1401 using MT as the female donor and as the recurrent parent in the further backcrosses necessary for introgression (Figure **1**). The process of introgression was based on visual screening on stereoscope for the presence of a high density of type-IV trichomes on adult leaves (5^th^ leaf in the MT background; Vendemiatti *et al*., 2017) and followed the procedure previously described by Pino *et al*. (2010). Plants were grown in a greenhouse under natural day-length conditions (Lombardi-Crestana *et al*., 2012).

### Plant genetic transformation

Constructs containing the green fluorescent protein (GFP) reporter driven by *pSlAT2*, which directed the expression of GFP to the tip of trichomes types I and IV (Schilmiller *et al*., 2012), were kindly provided by Dr. Robert Last (Michigan State University, USA). The constructs were introduced into *Agrobacterium tumefaciens* LBA4404 and used to transform MT and MT-*Get* as described by Pino *et al*. (2010). Plants regenerated under kanamycin selection, were acclimated in a greenhouse, and cultivated as described above.

### Trichome counting and phenotyping

Identification and counting of trichomes were carried out as described by Vendemiatti *et al*. (2017). At least 8 individuals per genotype were sampled, and four different samples were analyzed per plant on each leaf surface. Photographs were taken using a Leica S8AP0 (Wetzlar, Germany) at 50x magnification, coupled to a Leica DFC295 camera (Wetzlar, Germany). Counting and length measurements of trichomes were performed on the images using the manufacturer’s analytical program (Leica Application Suite 4.0).

### Scanning electron microscopy

Leaf samples were fixed in Karnovsky solution for 24 hours at 4°C. The material was then washed twice with 0.05 M cacodylate solution for 10 minutes and fixed again in osmium tetroxide for 1 hour. Subsequently, the samples were washed with distilled water and dehydrated in a series of acetone baths. The dehydrated samples were submitted to drying to the critical point, and subsequently gold plated. The observations were performed on a LEO 435 VP scanning electron microscope (SEMTech Solutions, Massachusetts).

### Fluorescence microscopy

For trichome-specific expression of GFP under the *SlAT2* promoter, analyses were carried out under a Nikon SMZ18 stereoscope attached to a Nikon DS-RI1 digital camera. Excitation at 480 nm and a 505 nm emission filter detected fluorescence specifically from GFP. For chloroplast fluorescence detection, trichomes were observed under a Carl Zeiss Axioskop 2 microscope coupled to an Axiocam MRc Zeiss camera using 540/625 nm excitation/emission filter.

### Mapping-by-Sequencing Analysis

A segregating BC_7_F_2_ population composed of 315 plants from the cross MT-*Get* (BC_6_F_n_) x MT was phenotyped for the presence of type-IV trichomes on adult (5^th^) leaves according to the methodology described above. Plants were classified into two populations: plants with leaves bearing type-IV trichomes (*Get*-like) and those with virtually no type-IV trichomes observed (MT-like). Five leaf discs (7-mm diameter) from each plant were collected and pooled into two populations according to phenotype before extraction of genomic DNA using the method described by Fulton *et al*. (1995). The genomic DNA was further purified with the MasterPure kit (Lucigen, MC85200) and submitted to sequencing on a HiSeq PE150bp (Illumina) at Novogene (https://en.novogene.com) with 30X depth. Fastq files for each population were concatenated and submitted to quality check by FastQC on the Galaxy platform (https://usegalaxy.org). Mapping of reads against the cv. Micro-Tom v.1 genome reference sequence (Sol_mic assembly: http://gbf.toulouse.inra.fr/Genome) was carried out with the “Map with BWA for Illumina” (v.1.2.3) software on Galaxy, using default parameters (Li & Durbin, 2009) to generate “.sam” files. The variant calling against the cv. Micro-Tom assembly was performed with Samtools Mpileup (v.1.8) (Li *et al*., 2009) and BCFtools call (v.1.6) to generate VCF files. Further comparisons between *Get*-like variants (S1) against the MT-like population (S4) were performed with BCFtools isec (v.1.6) on the command line interface. The variant allelic frequencies were filtered per site relative to the MT genome sequence using the following parameters: i) total depth of reading (10<DP<100); ii) allele frequency (AF>=0.8), which is defined as the alternative depth of reading (AD) divided by the total depth of reading (DP); and iii) number of variants per 1 Mbp window >= 30 (Mascher *et al*., 2014; Garcia *et al*., 2016). The analysis pipeline can be visualized in Supporting Information Figure **S1**.

### Micro-Tom and S. galapagense allele genotyping

Genomic DNA was extracted using the protocol described by Fulton et al. (1995). The DNA quantity and quality were determined using agarose gel electrophoresis and NanoDrop One spectrophotometer (Thermo Fisher Scientific). The genotyping was performed using CAPS markers that discriminate *Solanum lycopersicum* cv. MT and *Solanum galapagense* alleles (Table **S1**). Each 12-μL PCR reaction contained 1.0 μL DNA, 1.2 μL Taq buffer (10x), 1.5 μL MgCl_2_ (25 nM), 0.2 μL dNTPs (10 mM), 0.4 μL each primer (10 pM), 0.1 μL Taq DNA polymerase (5U/μL - Thermo Fisher Scientific), and 7.2 μL distilled water. The PCR programs were developed according to the optimum annealing temperatures and amplicon sizes of each primer set. The digestion reactions (10μL) contained 4.0 μL PCR product, 1.0 μL enzyme buffer (10x), 0.2 μL restriction enzyme, and 4.8 μL water. The products were analyzed on 1.5% (w/v) agarose gels, using SYBR^®^ Gold Nucleic Acid Gel Stain (Invitrogen).

### Plant phenotyping

Forty-eight-day old plants were used for measuring the length of the main stem and the length of the secondary branches of the plant, and the branching index was calculated according to Morris *et al*. (2001). For fruit weight measurements, MT and MT-*Get* plants growing in 250-mL pots were hand-pollinated with their own pollen.

Many ovaries were pollinated, but after fruit set confirmation (five days after pollination), we performed selective fruit thinning to allow only five fruits to develop and ripe on each plant.

### Herbivory test with Bemisia tabaci

Seeds were sown in plastic trays using coconut fiber substrate and remained in greenhouse conditions until the transplant. The seedlings were then transplanted into 8-liter pots with substrate. Each pot received 5 plants of the same genotype. The plants were kept in a greenhouse until 23 days after transplanting. During the interval between transplanting and the beginning of inoculation, the plants received the appropriate cultural treatments and fertilization. After 23 days, the pots were randomly placed in a greenhouse chamber (7m x 15m) highly infested with a whitefly population (*Bemisia tabaci*), where the insects are bred and kept exclusively for tomato resistance tests. The pots remained there for 7 days to allow egg lying on the leaves of the plants. After the inoculation period, a vase of each genotype, each containing 5 plants, was randomly collected. From these plants, 30 leaflets were collected to perform the counting of hatched nymphs.

### Rhodamine-B assay for acylsugar staining

Leaflets of *S. galapagense* and MT-*Get* were submerged in a 0.1% aqueous solution of Rhodamine-B for one minute. Subsequently, the samples were gently immersed in distilled water four times (serially) to remove the excess dye. The images of stained trichomes were taken as described above.

### LC-MS/MS analysis of surface extracts

This experiment was carried out by the Glandular Trichomes and Isoprenoid Biosynthesis Research Group at the Leibniz Institute for Plant Biochemistry (Halle, Germany). Semipolar metabolites were extracted by placing two tomato leaflets (of the 5^th^ leaf) in a 2-mL reaction tube containing 1 mL of methanol. After vortexing, the samples for 1 min the supernatant was transferred to a new tube, centrifuged for 5 min at 18,000 *g* and filled in a glass vial. The analysis of the extracted metabolites was performed on a LC-MS/MS system composed of an Acquity UPLC (Waters GmbH, Eschborn, Germany) and a TripleTOF 5600 mass spectrometer (SCIEX, Toronto, Canada). For the separation of the analytes, 5 µL of extracts were injected into a Nucleoshell RP 18 column (2.7 µm x 150 mm x 2 mm, Macherey-Nagel GmbH, Düren, Germany). A solvent system composed of A: 0.3 mM ammonium formate acidified with formic acid at pH 3, and B: acetonitrile, with the following gradient was used: 0-2 min: isocratic 95% A, 2-19 min: linear from 95% to 5% A, 19-22 min: isocratic 5% A, 22-22.01 min: linear from 5% A to 95% A, 22.01-24 min: isocratic 95% A. The flow rate was set to 400 µL/min throughout and the column temperature was 40°C. Analyte ionization was performed by electrospray ionization in negative mode with the following parameters: gas 1 = 60 psi, gas 2 = 70 psi, curtain gas = 35 psi, temperature = 600°C and ion spray voltage floating= -4500 V. CID fragment spectra were generated in SWATH mode (Hopfgartner *et al*., 2012) with mass windows of 33 Da and rolling collision energies from -10 to -80 V with a collision energy spread of 15 V. The integration of the peak areas was performed by Multiquant (Version 2.0.2; SCIEX, Toronto, Canada).

### GC-MS Acyl sugar quantification

This experiment was carried out at the Laboratory of Biochemistry and Instrumental Analysis of the Department of Agroindustry, Food, and Nutrition (ESALQ-USP). Leaves of the adult vegetative phase (5^th^ leaf) were collected, and the extraction was conducted according to the methodology described by Leckie *et al*. (2013). The compounds were separated via GC-2010 gas chromatography (Shimadzu Corp., Kyoto, Japan) attached to a QP 2010 Plus mass spectrometer (Shimadzu Corp., Kyoto, Japan), using Helium as the charging gas. For the separation of acyl groups from the acyl-sugar molecules, hexane was injected into a DB-WAX apolar column (0.25 mm diameter, 30 m length, and 0.25 μm film thickness). The data obtained were analyzed using the software Lab Solutions-GC/MS version 2.5 (Shimadzu Corp., Kyoto, Japan). Compound identification was based on the retention time of chromatographic peaks and fragments of the mass spectrometer, which were compared to available standards and data libraries (Wiley® 8 and FFNSC 1.3). The identified compounds were quantified using a calibration curve derived from the peak areas of the standards.

### Gene expression analyses

The expression of key structural genes involved in the AS biosynthesis pathway was performed by real-time PCR in MT, MT-*Get*, and the wild species *S. galapagense*.

The comparison was performed only in genotypes harboring type-IV trichome in all leaves. Total RNA was isolated from leaf pools using the mirVana™ Isolation Kit (Ambion) according to the manufacturer’s instructions. The RNA was quantified on a NanoDrop One UV-Vis Spectrophotometer (Thermo Scientific), and the RNA integrity was examined by gel electrophoresis. The total RNA was treated with TURBO DNA-free™ Kit (Invitrogen™) and subsequently used for cDNA synthesis using the SUPERSCRIPT™ IV 1^st^ Strand Synthesis kit (Invitrogen) according to the manufacturer’s instructions. Quantitative RT-PCR (qPCR) reactions were conducted in a 10-µL total volume using a 2× GoTaq® qPCR Master Mix (Promega) and run on an ABI 7500 qPCR thermocycler (Applied Biosystems). The constitutive housekeeping genes *ACTIN* (Solyc04g011500) and *ELONGATION FACTOR 1 ALPHA* (*EF*-*1*, Solyc06g005060) were used as internal controls. We used three biological repetitions, each composed of 5 leaves, and three technical repetitions. The threshold cycle (C_T_) was determined automatically by the instrument, and fold changes for each gene of interest were calculated using the equation 2^-ΔΔCt^ (Livak & Schmittgen, 2001). The qPCR primer sequences used are listed in Supporting Information Table **S1**.

### Statistical analyses

The LC-MS data were converted to the Log10 function before analysis. The statistical comparisons with ANOVA and the Student’s *t*-test were performed using the SAS software. The remaining results were compared statistically by the Student’s *t*-test using GraphPad Prism version 7.00 for Mac (GraphPad Software, La Jolla California USA, www.graphpad.com).

## ACCESSION NUMBERS

DNA-Seq raw dataset used for mapping-by-sequencing was submitted to the NCBI Sequence Read Archive (BioProject # XXXX - *data currently under submission*).

## ACKNOWLEDGEMENTS

This work was supported by funding from the Agency for the Support and Evaluation of Graduate Education (CAPES, Brazil), the National Council for Scientific and Technological Development (CNPq, Brazil), and the São Paulo Research Foundation (FAPESP, Brazil). This work was partially supported by FAPESP grants 2015/50220-2 and 2018/05003-1. CAPES is thanked for scholarships granted to RT and EV. EV, MSP and MHV received scholarships from FAPESP (grants #2016/22323-4, 2016/17092-3, and 2016/05566-0, respectively). SMA and LEPP received fellowships from CNPq (grants # 307893/2016-2 and 306518/2018-0, respectively). We also thank Dr. Mohamed Zouine (INP-Toulouse, France) for making the cv. Micro-Tom genome sequence available, and Dr. Robert Last (Michigan State University, USA) for the *pSlAT2*::*GFP* construct. We also thank our technician, Cassia R. F. Figueiredo, NAP/MEPA (ESALQ/USP) for assistance with electronic microscopy, Francisco Vitti for greenhouse assistance, and the former students Ariadne F. L. de Sá and Frederico A. de Jesus for PCR primer design.

## SHORT LEGENDS FOR SUPPORTING INFORMATION

**Figure S1** Mapping-by-sequencing bioinformatic analysis pipeline

**Figure S2** Absence of green fluorescence in non-transgenic MT-*Get* type-IV trichomes, and fluorescence of type-IV trichomes in *S. galapagense* showing the presence of chloroplasts.

**Figure S3** Ontogenetic sequence of type-IV and –V trichomes on both leaf surfaces of Micro-Tom (MT) and MT-*Get*.

**Figure S4** Counting of others trichomes types in *Solanum galapagense*, and density (mm^-2^) of additional trichome types on both leaf surfaces of Micro-Tom (MT) and MT-*Get*.

**Figure S5** Density (mm^-2^) of types-IV and –V trichomes on abaxial surfaces of leaves from MT-*Get*, and sublines MT-*Get*01, MT-*Get*02 and MT-*Get*03.

**Figure S6** Comparative scheme of some phenotypical and genetic traits from Micro-Tom (MT), MT-*Get* and *Solanum galapagense*.

**Figure S7** Representative pictures and quantification of whitefly (*Bemisia tabaci*) nymph infestation on Micro-Tom and MT-*Get* leaves. Representative micrographs of *Solanum galapagense* droplets on type-IV trichomes and their absence in MT-*Get*. In the inserts, trichomes were dyed with Rhodamine-B, reveling AS exudation in *S. galapagense*’s type-IV trichomes.

**Figure S8** GC-MS profile comparison between *Solanum galapagense* LA1401, Micro-Tom (MT) and MT-*Get* regarding acyl groups content. Full mass spectrum scan showing the relative abundance of ions for acyl group peaks found on extracts analyzed by GC-MS.

**Figure S9** Relative transcript accumulation of genes coding for *ASH* enzymes, and ABC transporter on leaf tissues of MT-*Get* and *S. galapagense*.

**Table S1** Oligonucleotide sequences used for molecular characterization.

**Table S2** Genomic coordinates of the genetic variation from *S. galapagense* present at high frequencies (≥0.8) in the *Get*-like phenotypical group of the MT-*Get* segregating population.

**Table S3** GC-MS profiles of acyl groups in *S. galapagense*, MT-*Get* and Micro-Tom (MT).

**Data S1** Genetic polymorphism found in *Get*-introgressed populations through DNA- sequencing analysis.

## SUPPORTING INFORMATION LEGENDS

**Figure S1** Mapping-by-sequencing bioinformatics analysis pipeline.

**Figure S2** Absence of green fluorescence in the tip cells of non-transgenic MT-*Get* type-IV trichomes as seen in light (a) and fluorescent (b) microscopy. (c) Fluorescence of type-IV trichomes showing chloroplast (red color) in *S. galapagense*. Scale bar=20 μm.

**Figure S3** Quantification of type-IV and –V trichomes in the adaxial (a-b) and abaxial (c-d) surfaces of cotyledons (Cot), first (L1), second (L2), third (L3), fourth (L4), fifth (L5) and sixth (L6) leaves of Micro-Tom (black bars) and MT-*Get* (white bars). Data are mean (n = 30) ± SEM. Asterisks indicate mean significantly different from the control MT, according to Student *t*-test *P*< 0.001 (***).

**Figure S4** (a) Density (mm^-2^) of types-I and –IV trichomes in both surfaces of leaves from *S. galapagense*. (b, c) Density (mm^-2^) of others trichomes types in adaxial (b) and abaxial surfaces (c) of MT and MT-*Get*. Data are mean (n = 35) ± SEM. Asterisks indicate significant differences when compared with reference sample according to Student’s *t-*test *P* < 0.001 (^***^).

**Figure S5** Density (mm^-2^) of types-IV and –V trichomes in abaxial surfaces of 5^th^ leaves from MT-*Get* (a, b) and their derived sublines MT-*Get*01 (c, d), MT-*Get*02 (e, f) and MT-*Get*03 (g, h). MT-*Get*01, 02 and 03 are BC_7_F_n_ lines harbouring *S. galapagense*’s chromosome 1, 2 and 3 segments. Data are mean (n=30) ± SEM.

**Figure S6** (a) Phenotype of representative 35-day-old Micro-Tom (MT) and the MT-*Get* plants. Scale bar=5 cm. (b) Branching Index values (n = 15). (**c**) Main stem height (n = 15). (d) Average fruit weight of MT-*Get* (n = 15). Fruits and sepals from MT (e), MT-*Get* (f), and *S. galapagense* (g). Note the small calyx and the slightly smaller fruit of the MT-*Get*, which are, respectively, the expected effects of the *EJ-2* and *FW3*.*2* alleles from *S. galapagense*. Note that MT-*Get* has the same red fruit that is characteristic of MT due to an absence of the *S. galapagense B* allele (see Fig. **5**). The presence of the B allele in *S. galapagense* produces an orange fruit (g) (Ronen *et al*., 2000). Scale bar =5 mm. (h) PCR-based markers showing the presence of the *S. galapagense* alleles *EJ-2* and *FW3*.*2* (see Fig. **5**). The asterisks indicate significant statistical differences according to the Student’s *t*-test at *P*< 0.05 (*) or *P*< 0.001 (^***^).

**Figure S7** Representative photographs of whitefly (*Bemisia tabaci*) nymph infestation on Micro-Tom (a) and MT-*Get* (b) leaves. Scale bar=250 μm. (c) Quantification of *Bemisia tabaci* nymphs in MT-*Get* compared to the control MT (n=30). The data are not statistically different according to the Student’s *t*-test (*P* <0.05). (d, e) Representative micrographs of *Solanum galapagense* droplets on type-IV trichomes (d) and their absence in MT-*Get* (e). In the inserts, trichomes were dyed with Rhodamine-B, reveling AS exudation in *S. galapagense* type-IV trichomes.

**Figure S8** (a) GC-MS comparison between *Solanum galapagense* LA1401, Micro-Tom and MT-*Get* regarding acyl groups content. The scale at the y-axis is in arbitrary units. (b) Full scan mass spectrum showing relative abundance of ions for acyl groups peaks found on extracts analysed by GC-MS.

**Figure S9** Relative transcript accumulation of the (a-c) *ASHs* enzymes and (d) ABC transporter in leaves of MT-*Get* and *S. galapagense*. qRT-PCR values are means ± SE of three biological samples. Asterisks indicate a significant difference when compared with reference sample according to Student’s *t*-test *P* < 0.05 (^*^); *P* < 0.01 (^**^); *P* < 0.001 (^***^).

**Table S1** Oligonucleotide sequences used in this work.

**Table S3** GC-MS content of acyl groups in *S. galapagense*, Micro-Tom (MT) and MT-*Get* leaves.

