## Supplemental Information for "Introgression of type-IV glandular trichomes from *Solanum galapagense* to cultivated tomato reveals genetic complexity for the development of acylsugar-based insect resistance"

### Supporting Information

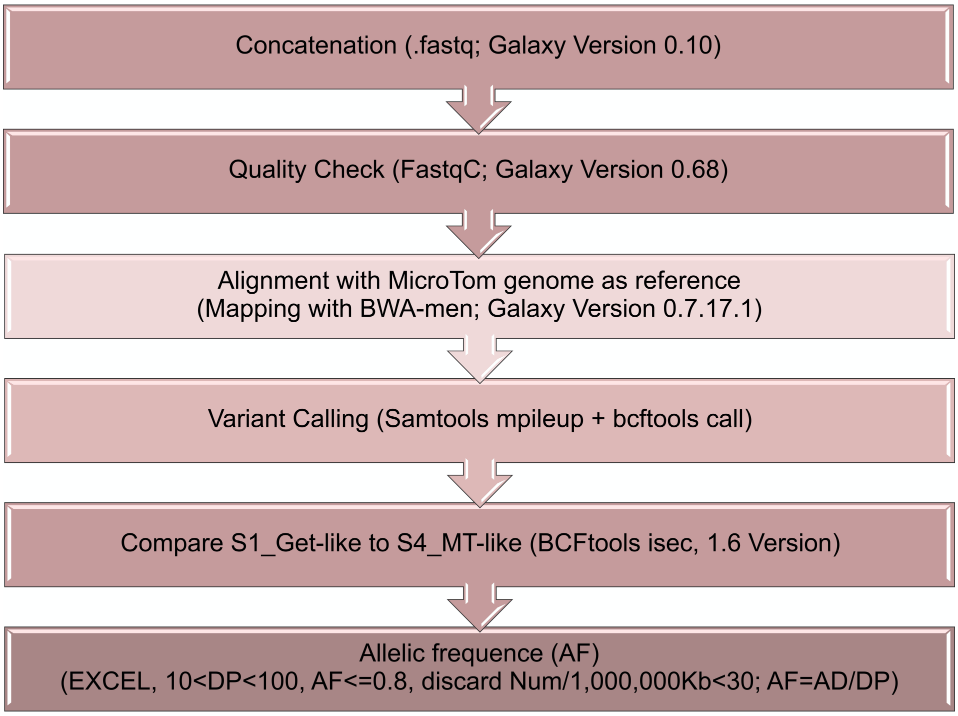


**Figure S1** Mapping-by-sequencing bioinformatics analysis pipeline.

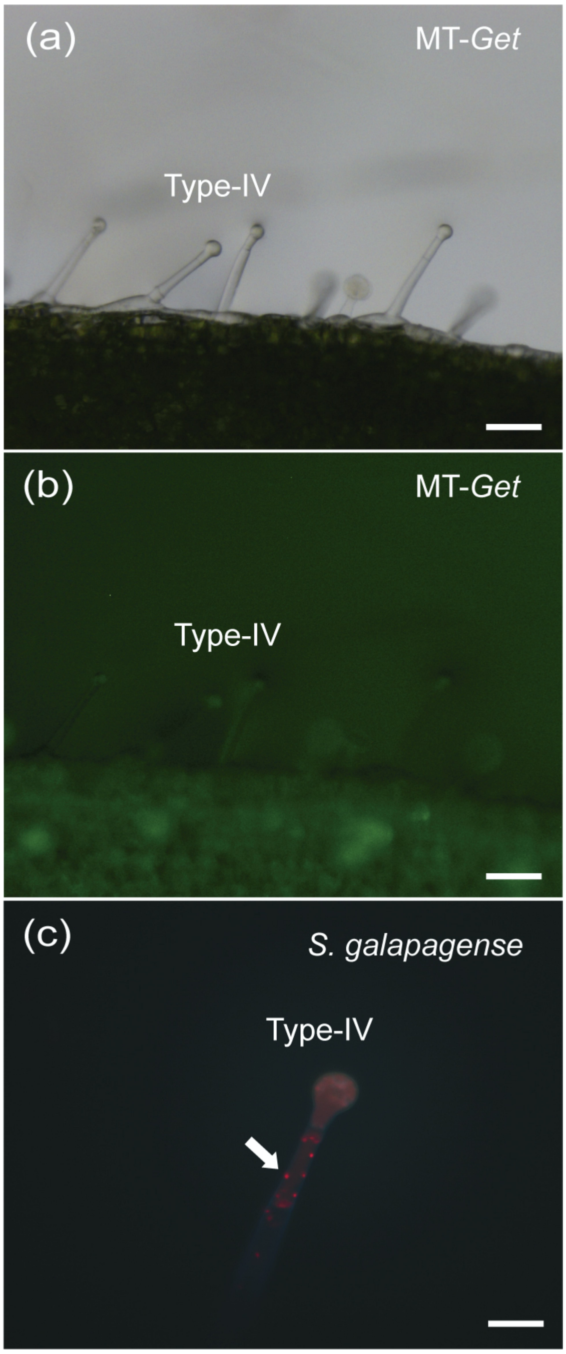


**Figure S2** Absence of green fluorescence in the tip cells of non-transgenic MT-*Get* type-IV trichomes as seen in light (a) and fluorescent (b) microscopy. (c) Fluorescence of type-IV trichomes showing chloroplast (red color) in *S. galapagense*. Scale bar=20 μm.

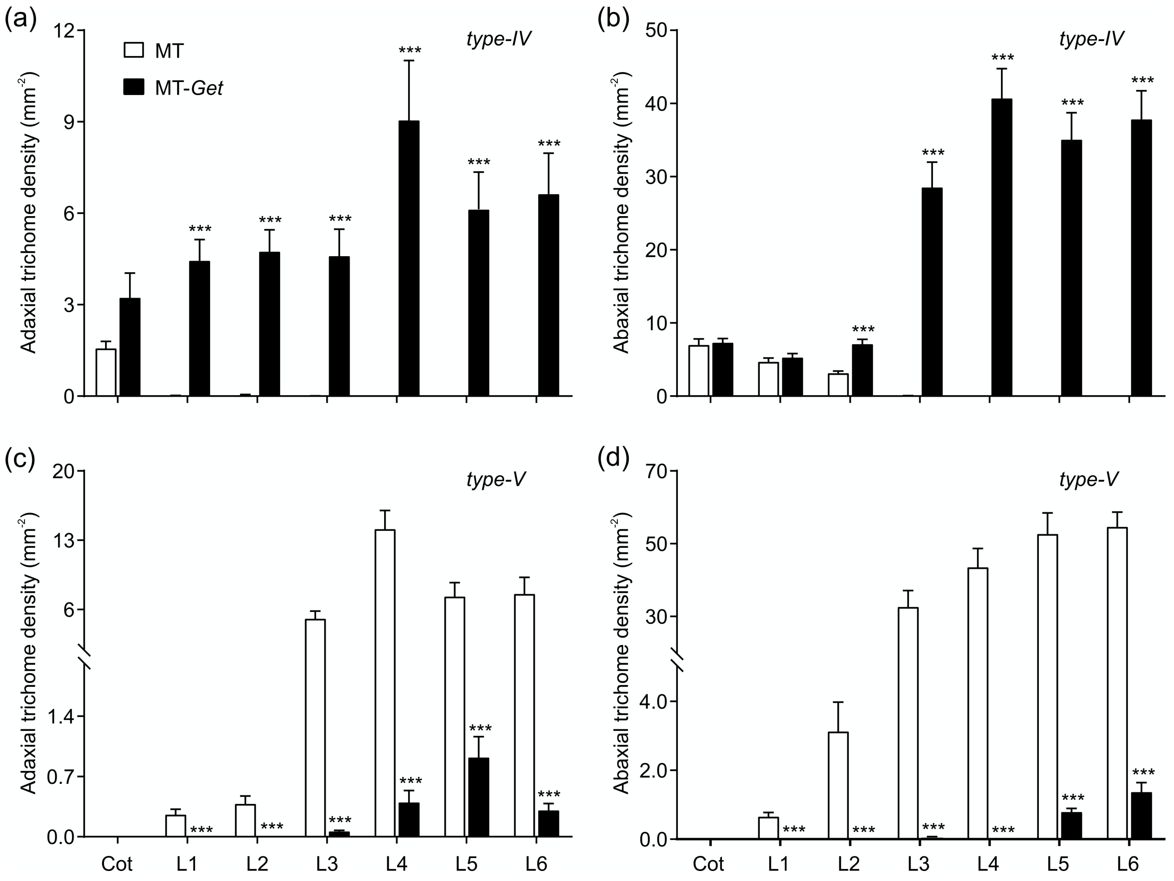


**Figure S3** Quantification of type-IV and –V trichomes in the adaxial (a-b) and abaxial (c-d) surfaces of cotyledons (Cot), first (L1), second (L2), third (L3), fourth (L4), fifth (L5) and sixth (L6) leaves of Micro-Tom (black bars) and MT-*Get* (white bars). Data are mean (n = 30) ± SEM. Asterisks indicate mean significantly different from the control MT, according to Student *t*-test $P< 0.001$ (***).

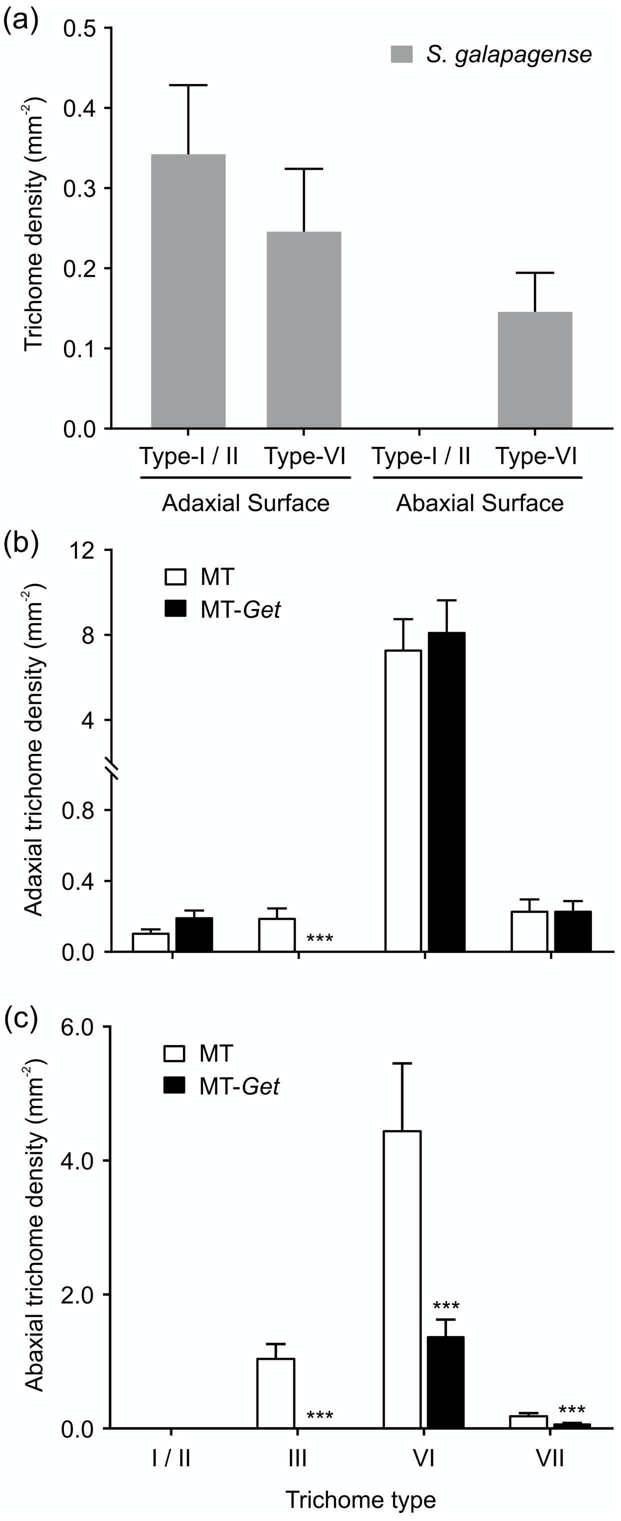


**Figure S4** (a) Density ($mm^{-2}$) of types-I and –IV trichomes in both surfaces of leaves from *S. galapagense*. (b, c) Density ($mm^{-2}$) of others trichomes types in adaxial (b) and abaxial surfaces (c) of MT and MT-*Get*. Data are mean (n = 35) ± SEM. Asterisks indicate significant differences when compared with reference sample according to Student's *t*-test $P < 0.001$ (***).

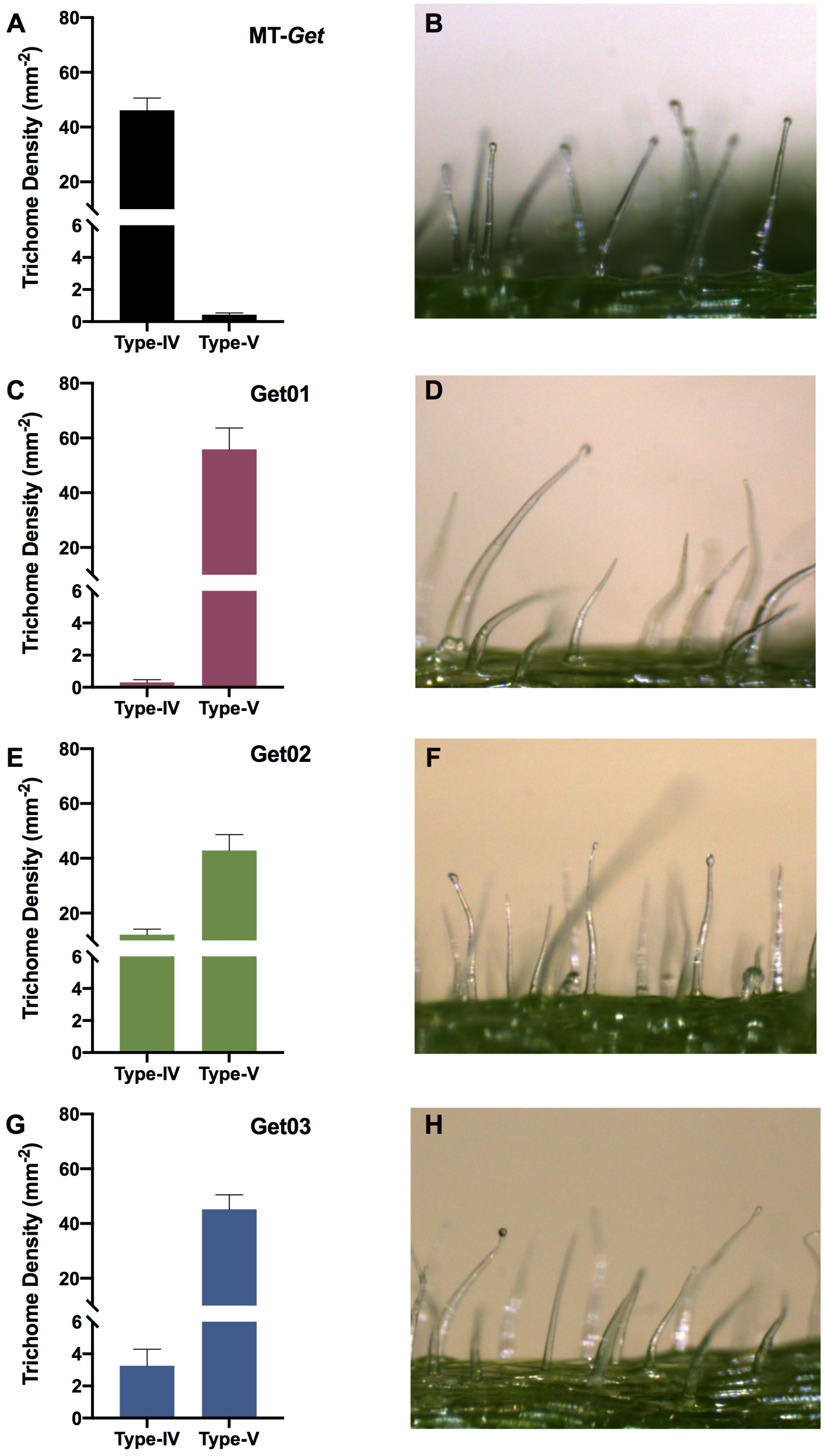


**Figure S5** Density (mm$^{-2}$) of types-IV and –V trichomes in abaxial surfaces of 5$^{th}$ leaves from MT-*Get* (a, b) and their derived sublines MT-*Get*01 (c, d), MT-*Get*02 (e, f) and MT-*Get*03 (g, h). MT-*Get*01, 02 and 03 are $BC_7F_n$ lines harbouring *S. galapagense*´s chromosome 1, 2 and 3 segments. Data are mean (n=30) ± SEM.

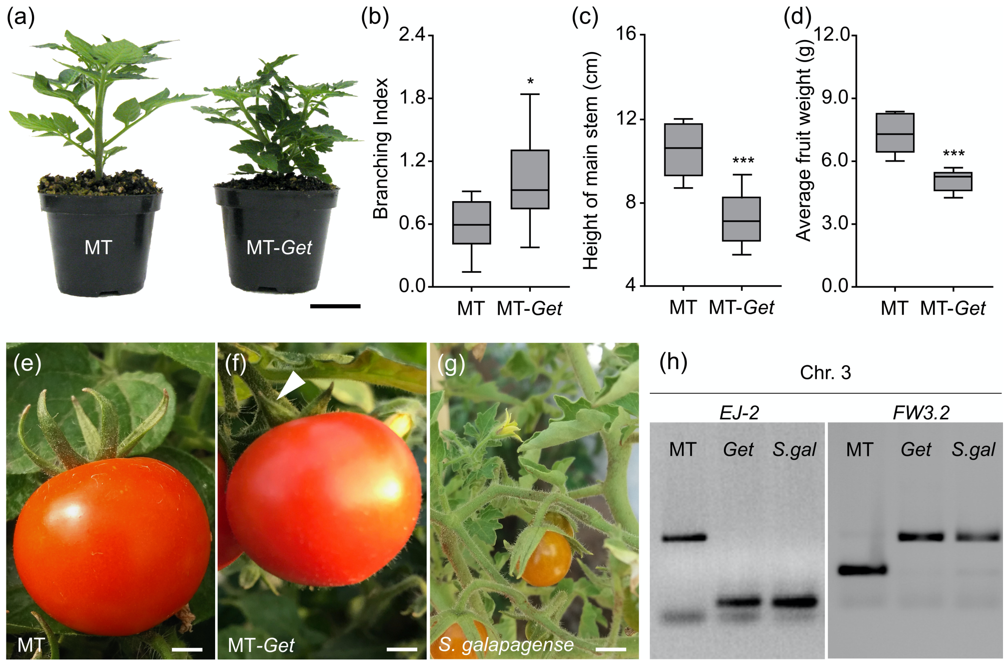


**Figure S6** (a) Phenotype of representative 35-day-old Micro-Tom (MT) and the MT-*Get* plants. Scale bar=5 cm. (b) Branching Index values (n = 15). (**c**) Main stem height (n = 15). (d) Average fruit weight of MT-*Get* (n = 15). Fruits and sepals from MT (e), MT-*Get* (f), and *S. galapagense* (g). Note the small calyx and the slightly smaller fruit of the MT-*Get*, which are, respectively, the expected effects of the *ej-2* and *fw3.2* alleles from *S. galapagense*. Note that MT-*Get* has the same red fruit that is characteristic of MT due to an absence of the *S. galapagense*´s *B* allele (see Figure **5**). The presence of the B allele in *S. galapagense* produces an orange fruit (g) (Ronen *et al.,* 2000). Scale bar =5 mm. (h) PCR-based markers showing the presence of the *S. galapagense* alleles *EJ-2* and *FW3.2* (see Figure 5). The asterisks indicate significant statistical differences according to the Student's *t*-test at *P*< 0.05 (*) or *P*< 0.001 (***).

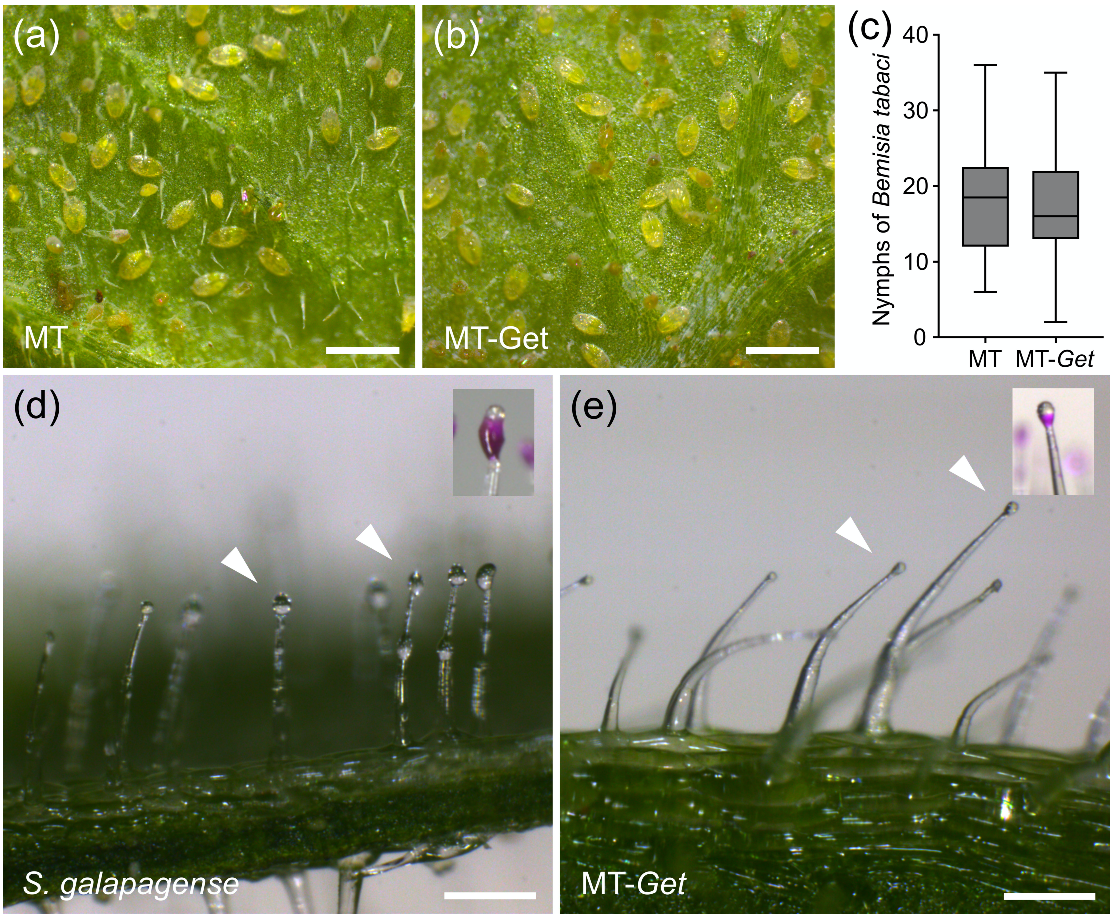


**Figure S7** Representative photographs of whitefly (*Bemisia tabaci*) nymph infestation on Micro-Tom (a) and MT-*Get* (b) leaves. Scale bar=250 µm. (c) Quantification of *Bemisia tabaci* nymphs in MT-*Get* compared to the control MT (n=30). The data are not statistically different according to the Student's *t*-test (*P* <0.05). (d, e) Representative micrographs of *Solanum galapagense* droplets on type-IV trichomes (d) and their absence in MT-*Get* (e). In the inserts, trichomes were dyed with Rhodamine-B, reveling AS exudation in *S. galapagense*´s type-IV trichomes.

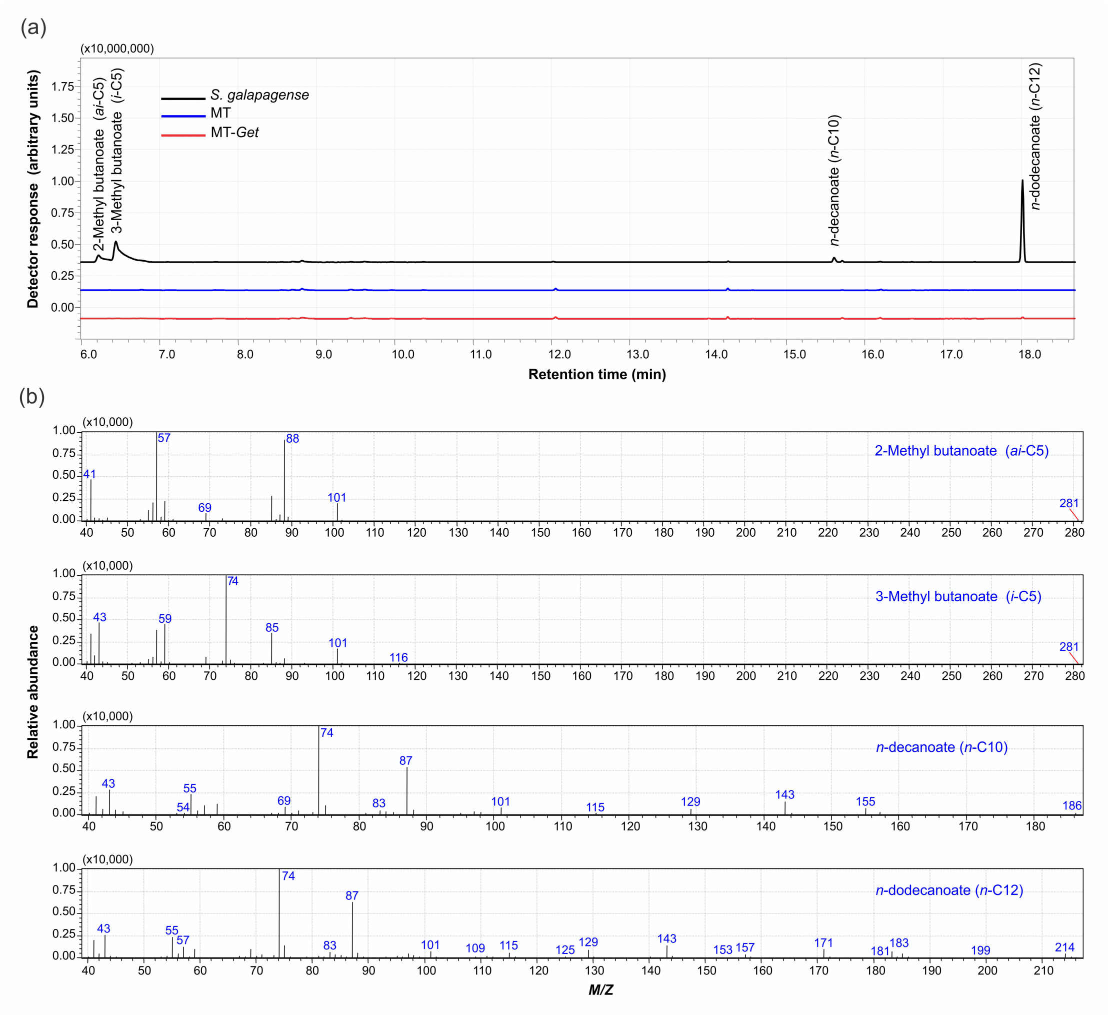


**Figure S8** (a) GC-MS comparison between *Solanum galapagense* LA1401, Micro-Tom and MT-*Get* regarding acyl groups content. The scale at the y-axis is in arbitrary units. (b) Full scan mass spectrum showing relative abundance of ions for acyl groups peaks found on extracts analysed by GC-MS.

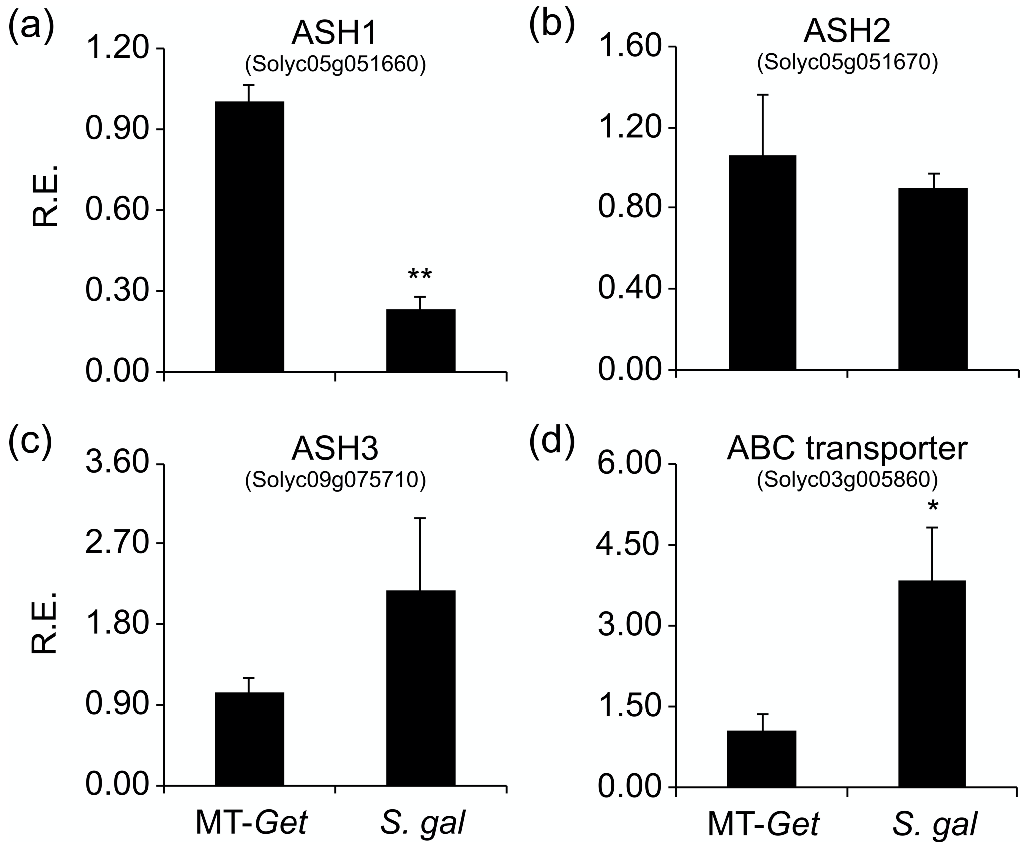


**Figure S9** Relative transcript accumulation of the (a-c) *ASHs* enzymes and (d) ABC transporter in leaves of MT-*Get* and *S. galapagense*. qRT-PCR values are means ± SE of three biological samples. Asterisks indicate a significant difference when compared with reference sample according to Student's *t*-test $P < 0.05$ (*); $P < 0.01$ (**); $P < 0.001$ (***).

**Table S1.** Oligonucleotide sequences used in this work.

| Primer | Sequence | Access | Utilization | Reference |
|---|---|---|---|---|
| 1B | AGCACTTATGGGCCTGTTCA<br>CAGCGCATCTTGAGTGTTGT | Solyc01g088100 | CAPS marker | This work |
| 2A | ACAACGACTAGAACGGCTGG<br>CCCATCACAACCTAATGTCTAAGG | Solyc02g092570 | CAPS marker | This work |
| 2B | GCACAACATCCACCCTTCTT<br>CAAGCACATACGATTGCAGA | Solyc02g093030 | CAPS marker | This work |
| 2C | GAAGGGTCGGGTACAAGGTAA<br>CCACTCTCTTAGCCTGAATGG | Solyc02g094500 | CAPS marker | This work |
| 3A | GCTCTGAAGATCCTACACATGAAA<br>TCCCATGCTCTTTCTGGAAC | Solyc03g019800 | CAPS marker | This work |
| 3B | CGGCGTAAGACAAACACGTTC<br>GCATTATCAACTCATCTACCAGTCC | Solyc03g083830 | CAPS marker | This work |
| 3C | TTGGGAGATTAGCGTAAGGAG<br>ATAGGAGCAACAAGGTACATGC | Solyc03g093610 | CAPS marker | This work |
| 6A | GCAGTGTGATGCTGGTGTTT<br>AGGCGACTTGGTTGTCACTC | Solyc06g005060 | CAPS marker | This work |
| 6B | TGTTGTTTCCCTTGTTGCAG<br>GCCAGTTTGAACCAGACCAG | Solyc06g009730 | CAPS marker | This work |
| 6C | AGCTAAGATGGAGGGAGATGC<br>CCCGAGGTGTAAACTGTAACG | Solyc06g048460 | CAPS marker | This work |
| 6D | AAAACACGGCGATGAGAAAT<br>TTTCCTCTCCAATTCCCAAT | Solyc06g060210 | CAPS marker | This work |
| *FW2.2* | GGTGGTGTTGATGTGGAGTGAGTG<br>GGCAGATACATAGTGAGGAGGAAC | Solyc02g090730 | CAPS marker | This work |
| *DWARF* | TGATCCATATTCGTTCAATCCA<br>CGTGATTATGTTAGCGGGAAT | Solyc02g089730 | CAPS marker | This work |

| | | | | |
|---|---|---|---|---|
| *FW3.2* | GAGATAACGGGTTAAATAGAGT<br>TAGTTAGGATAGTTATAGTTTGC | Solyc03g114940 | CAPS marker | This work |
| *EJ-2* | CACAATTCATGCTGGATCAGC<br>CGGAGTAATCTATTAGATTCTGC | Solyc03g114840 | CAPS marker | This work |
| *SP5G* | TAATTACGCAGTGACGAAGCA<br>TTGACACAGAGTTCGAGAACG | Solyc05g053850 | CAPS marker | This work |
| *SP* | GGGTTGAAGTTCATGGTGGT<br>AGTGCCTGGAATGTCTGTGA | Solyc06g074350 | CAPS marker | This work |
| *PTS* | AGGAAGTGATTGACCCATGC<br>CCCCAAACACCACTATCTAAGC | Solyc06g072480 | CAPS marker | This work |
| *ASAT1* | GGGAGGCCAAGACAAGTTGATA<br>TGGAGAAGCAAACTGAAGAAAATC | Solyc12g006330 | qPCR | Fan et al., 2016 |
| *ASAT2* | GACTCCATTCGTCCATCTTTACTTC<br>TTTGACTTCTTCCTTCTCCTTTCTTA | Solyc04g012020 | qPCR | |
| *ASAT3* | TTTCTTCCCTTTACCGTCTGAA<br>TGAACAAGTGCTGAGGCAAC | Solyc11g067270 | qPCR | This work |
| *ASAT4* | GGTGGTCGTGATGTCCCTAA<br>GCCCTCCTTGTTAGCAGTTG | Solyc01g105580 | qPCR | This work |
| *ASH1* | TCTTTCATCCAACGTGATTAACATTT<br>ATCTACACCACAGAACACTACCAATA | Solyc05g051660 | qPCR | This work |
| *ASH2* | GCGACCCACTGAATAGCATC<br>GGCGGAGGATTTGTATTAGGA | Solyc05g051670 | qPCR | This work |
| *ASH3* | CTACACTCAAATCAACTTCCATACCATACA<br>ATAGAACCGTTCGACTCGACCATTT | Solyc09g075710 | qPCR | Schilmiller *et al.*, 2016 |
| ABC transporter | TCCGAAGGGATGATGGAG<br>GCAGAAGACCAAATACAGGGTAA | Solyc03g005860 | qPCR | This work |
| *ACTIN* | GGTCCCTCTATTGTCCACAG<br>TGCATCTCTGGTCCAGTAGGA | Solyc04g011500 | qPCR | Silva *et al.*, 2018 |

| | | | | |
|---|---|---|---|---|
| *EF1α* | AAGCCCATGGTTGTTGAGAC<br>TTCTTGACAACACCCACAGC | Solyc06g005060 | qPCR | Pinto *et al*., 2017 |

**Table S2.** Genomic coordinates of the genetic variation from *S. galapagense* present at high frequencies (≥0.8) in the *Get*-like phenotypical group of the MT-*Get* segregating population.

| Chromosome | Corresponding Region (bp) |
|---|---|
| 1 | 82,558,349 – 83,597,817 |
| 2 | 53,604,856 - 55,297,272 |
| 3 | 61,495,951 - 65,007,167 |
| 5 | 19,540,000 - 19,637,400 |
| 6 | 3,688 - 38,647,717 |

**Table S3.** GC-MS content of acyl groups in *S. galapagense*, MT-*Get* and Micro-Tom (MT).

| RT | Acyl Groups Content | Genotypes | | |
|---|---|---|---|---|
| | | ***S. galapagense*** | **MT-*Get*** | **MT** |
| 1 | 2-Methyl Butanoate | 0.77 ± 0.30 | 0.00 | 0.00 |
| 2 | 3-Methyl Butanoate | 2.86 ± 1.20 | 0.00 | 0.00 |
| 3 | Methyl Decanoate | 0.18 ± 0.07 | 0.00 | 0.00 |
| 4 | Methyl Dodecanoate | 2.92 ± 0.90 | 0.06 ± 0.00 | 0.00 |

Data are mean (n =8) ± SD.
